# An immune-poised state of human bronchial epithelial cells mediates RSV resistance in adults

**DOI:** 10.64898/2026.02.02.703395

**Authors:** Caiqi Zhao, Yan Bai, Hongmei Mou, Paul H. Lerou, Daisy W. Leung, Harish Narasimhan, Xiaoqin Wei, Jie Sun, Jin-Fu Xu, Rachel Fearns, Xingbin Ai

**Affiliations:** Division of Newborn Medicine, Department of Pediatrics, Massachusetts General Hospital, Boston, MA 02114, USA; The Mucosal Immunology and Biology Research Center, Massachusetts General Hospital for Children, Boston, MA 02114, USA; Department of Medicine, Washington University School of Medicine, St. Louis, MO 63110, USA; Beirne B. Carter Center for Immunology Research, University of Virginia, Charlottesville, VA 22904, USA; Department of Respiratory and Critical Care Medicine, Tongji Hospital, School of Medicine, Tongji University, Shanghai 200065, China; Institute of Respiratory Medicine, School of Medicine, Tongji University, Shanghai 200065, China; Department of Virology, Immunology & Microbiology, Boston University Chobanian & Avedisian School of Medicine, Boston, MA 02118, USA

**Keywords:** age, basal stem cells, ciliated cells, respiratory syncytial virus, NS2, RIG-I, STAT3, H3K27me3

## Abstract

Respiratory syncytial virus (RSV) impairs STAT3 activation in infected ciliated cells in infants, but not adults, causing apoptosis to facilitate virus spread. How age regulates the STAT3 response to RSV in human bronchial epithelial cells (BEpiCs) is unknown. Here, we identify an age-associated immune-poised physiological state in human bronchial basal stem cells (BSCs) and progeny BEpiCs, rendering robust RIG-I to guard against the inactivation of RIG-I-STAT3 signaling by RSV nonstructural protein 2. In contrast, infant BEpiCs are vulnerable to such inactivation by RSV due to lower RIG-I levels. We show that acute respiratory viral infection in specific pathogen-free mice, rather than physical age, triggered sustained epigenetic and transcriptomic remodeling in BEpiCs that resembled age-associated changes in human BEpiCs. In addition, cytokine pretreatment of human neonatal BSCs induced an adult-like immune-poised state that was maintained in progeny BEpiCs to protect against RSV. Therefore, respiratory infection and inflammation during a lifetime imparts the effect of age to BSCs, which fortifies RIG-I in progeny cells to mediate RSV resistance.

---

RSV primarily targets bronchial epithelial ciliated cells^1,2^ and is the most common cause of bronchiolitis in children younger than 2 years of age, causing respiratory distress and even death^3–10^. RSV infection in young children also increase susceptibility to asthma later in life^11–14^. Prophylactic strategies reduce medically attended RSV disease in infants^15–17^. However, for infants with active RSV infection, supportive care remains the only treatment option. In contrast, RSV-infected adults (<65 years of age) often present cold-like symptoms. How age regulates susceptibility to severe RSV infection in infants is not fully understood partially due to difficulties of lung biopsy and lack of clinically-relevant experimental models^18,19^.

We have generated age-appropriate human BEpiCs in air-liquid interface (ALI) cultures of BSCs derived from tracheal aspirates (TA)^20–23^ to model RSV infection^24,25^. Since BSCs derived from TA and lung biopsy samples have comparable transcriptome profiles and differentiation potentials^22,23^, our TA approach enables BSC derivation from young children that otherwise is contraindicated. Age has no effect on RSV entry, primary viral replication, or the interferon (IFN) response at the early stage of infection in human BEpiCs^24^. However, compared to RSV-infected adult ciliated cells that robustly activate STAT3 (p-STAT3^Y705^) above an already higher baseline to prevent apoptosis, infant ciliated cells are impaired in the p-STAT3^Y705^ response to RSV, rendering apoptotic cell extrusion to facilitate virus spread^24^. Notably, SARS-CoV-2, which also targets ciliated cells^26,27^, exhibits similar infection between neonatal and adult BEpiCs^24^. Therefore, RSV-specific interaction with ciliated cells is regulated by age to determine the severity of infection.

RSV deploys nonstructural protein 1 (NS1) and NS2 to interact with host factors and suppress interferon (IFN) signaling and innate immune responses^28–31^. RSV NS1 and NS2 share little sequence homology and are unique among Mononegavirales viruses^32^. The helix α1 structure of NS2 specifically binds to the N-terminal caspase activation and recruitment domains (CARDs) of RIG-I and MDA5, which prevents conformational changes required for the activation of these two viral RNA sensors^33^. RIG-I and MDA5 show cell context-dependent activities^34^. RIG-I plays a major role in the antiviral response in human nasopharyngeal epithelial cells (upper airway) but not BEpiCs (lower airway)^35^. In addition, RIG-I, but not MDA5, is required to trigger spleen tyrosine kinase (Syk)-mediated p-STAT3^Y705^ to promote defense again viruses^36^. Whether RSV NS2-specific interaction with RIG-I regulates the p-STAT3^Y705^ response to RSV in human BEpiCs and if so, whether age affects the interaction are unknown.

Here, we investigate the molecular underpinning of age-associated RSV-BEpiC interaction in humans. Our findings uncover an underappreciated role of respiratory infection/inflammation during a lifetime in fostering an immune-poised state in adult BSCs and progeny cells, which mediates protection against severe RSV infection through the RSV NS2-RIG-I-STAT3 axis.

## Results

### Human BEpiCs exhibit an immune-poised physiological state with age

We assessed age-associated changes in the transcriptomes of ciliated cells using single cell RNA-sequencing (scRNA-seq) datasets of healthy donor lungs from 13 young children (<2 years of age) and 12 adults (29-56 years of age) (available via LungMap) (Extended data Fig. 1). Gene set enrichment assay (GSEA) found 15 most significantly enriched hallmark pathways with age in adult ciliated cells, including inflammatory responses to TNFα, IFN, and IL6 among the top 6 pathways (Fig. 1a and Extended data Fig. 1). BSCs differentiate into ciliated cells during homeostasis and regeneration of human lungs^37^. To test whether BSCs were the cell origin of the age-associated gene signature in ciliated cells, we complemented scRNA-seq with bulk RNA-seq of neonatal and adult, BSCs and ALI cultures^22–24^ to overcome limited capture efficiency and few BSCs from young children in the scRNA-seq datasets. We found 11 out of the top 15 pathways similarly enriched by age in BSCs and ALI cultures as in ciliated cells, including the 5 inflammatory response pathways (Fig. 1a). In contrast, age had no such effect on AT1 or AT2 cells. Therefore, age is associated with the enrichments in inflammatory response pathways in BSCs and progeny BEpiCs *in vivo* and *in vitro*.

**Fig. 1.**
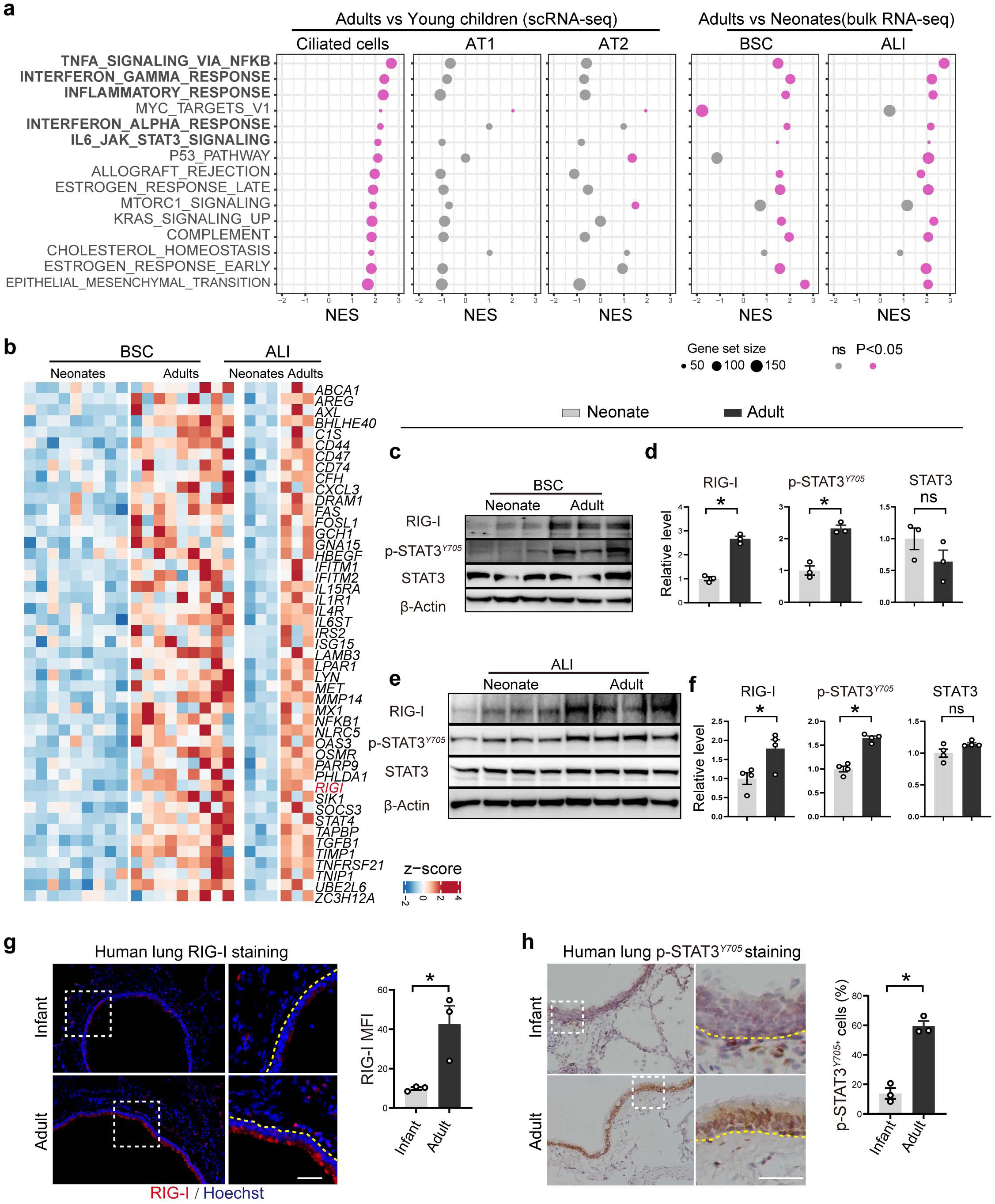
Human BEpiCs exhibit an immune-poised physiological state with age. **a**, Dot plots for the 15 most significantly enriched hallmark pathways with age in ciliated cells in donor lungs of young children (N=13) and adults (N=12) by scRNA-seq dataset analyses. The status of age association of these 15 pathways in AT1 and AT2 cells, neonatal and adult BSCs (N=9 per age group) and day 21 ALI cultures (N=3 per age group) was also evaluated. **b**, Heatmap of the 47 age-associated inflammatory response genes in BSCs and ALI cultures. Each lane represents one donor. **c**,**e**, Representative Western blot assays of RIG-I, p-STAT3^Y705^, and STAT3 in neonatal and adult BSCs and ALI cultures. β-Actin was loading control. Each lane is one donor. **d**,**f**, Quantification of Western blot results by densitometry normalized to β-Actin. **g**,**h**, Representative antibody staining for RIG-I (fluorescence) and p-STAT3^Y705^ (chromogenic) in infant and adult lungs (N=3 donors per age). Cells residing in lung mesenchyme in the same images showed comparable levels of staining for RIG-I and p-STAT3^Y705^ between the two ages. Nuclei were counter stained by Hoechst dye in **g** and hematoxylin in **h**. Data were quantified by mean fluorescence intensity (MFI) for RIG-I staining and the percentage of bronchial epithelial cells positive for p-STAT3^Y705^. Panels on the right show enlarged outlined areas in panels on the left. The yellow dotted line marks basement membrane. *p<0.05 and ns, not significant by Student’s t-test (two-tailed).

Among ∼500 inflammatory response genes, 56 genes were differentially expressed by more than 30% (adjusted p value <0.05) between neonates and adults in both BSCs and ALI cultures, including 47 upregulated and 0 downregulated genes (Fig. 1b and Extended Data Fig. 2). Only a small number of genes showed a more than 2-fold difference in expression between neonates and adults, which likely reflects physiological changes in human BEpiCs with age *in vivo*^38^. A subset of the 47 genes were reliably detected in more than 20% of ciliated cells by scRNA-seq and all were upregulated in adults compared to young children (Extended data Fig. 2). RIG-I was the only viral RNA sensor found upregulated with age by bulk RNA-seq, which was validated at the protein level by Western blot (Fig. 1b-f). Antibody staining consistently showed more robust RIG-I expression in adults than infants throughout the human bronchial epithelium in ALI cultures and donor lungs, including ciliated cells (Fig. 1g and Extended Data Fig. 3). A few cells in the lung mesenchyme had comparable staining for RIG-I and p-STAT3^Y705^ between the two age groups (Fig. 1g,h), providing additional evidence for a BEpiC-specific effect of age on the inflammatory response pathways. In parallel, baseline p-STAT3^Y705^ was elevated with age (Fig. 1c-f,h) as previously reported^24^. Taken together, age is associated with elevated levels of the inflammatory response signature and baseline p-STAT3^Y705^ in BEpiCs, reflecting an immune-poised physiological state of the bronchial epithelial barrier in adults.

### RSV NS2 induces severe infection and inactivates the p-STAT3^Y705^ response in neonates

We tested the role of NS1 and NS2 in severe RSV infection in neonatal BEpiCs in ALI using a doxycycline (Dox)-inducible lentiviral shRNA approach^24^ with a knockdown efficacy of 40-60% (Fig. 2a,c,d). This approach circumvents technical variables caused by attenuated growth of mutant RSV with deletions of NS1 or NS2^29,39,40^. NS1 or NS2 knockdown had no effect on the number of RSV-infected cells or viral gene expression at 1-day post-infection (dpi) (Fig. 2b-d). Only NS1 knockdown doubled the mRNA level of *IFNL1* and *IFNL3* (Fig. 2c,d), consistent with NS1 being a more robust repressor of the host IFN response than NS2^41^. At 2 dpi, despite an earlier anti-viral IFN response, NS1 knockdown had no effect on severe RSV infection in neonatal BEpiCs (Fig. 2b,e-g). In contrast, NS2 knockdown significantly reduced apoptosis of RSV-infected ciliated cells and virus spread. Therefore, NS2 is required for severe RSV infection in neonatal BEpiCs.

**Fig. 2.**
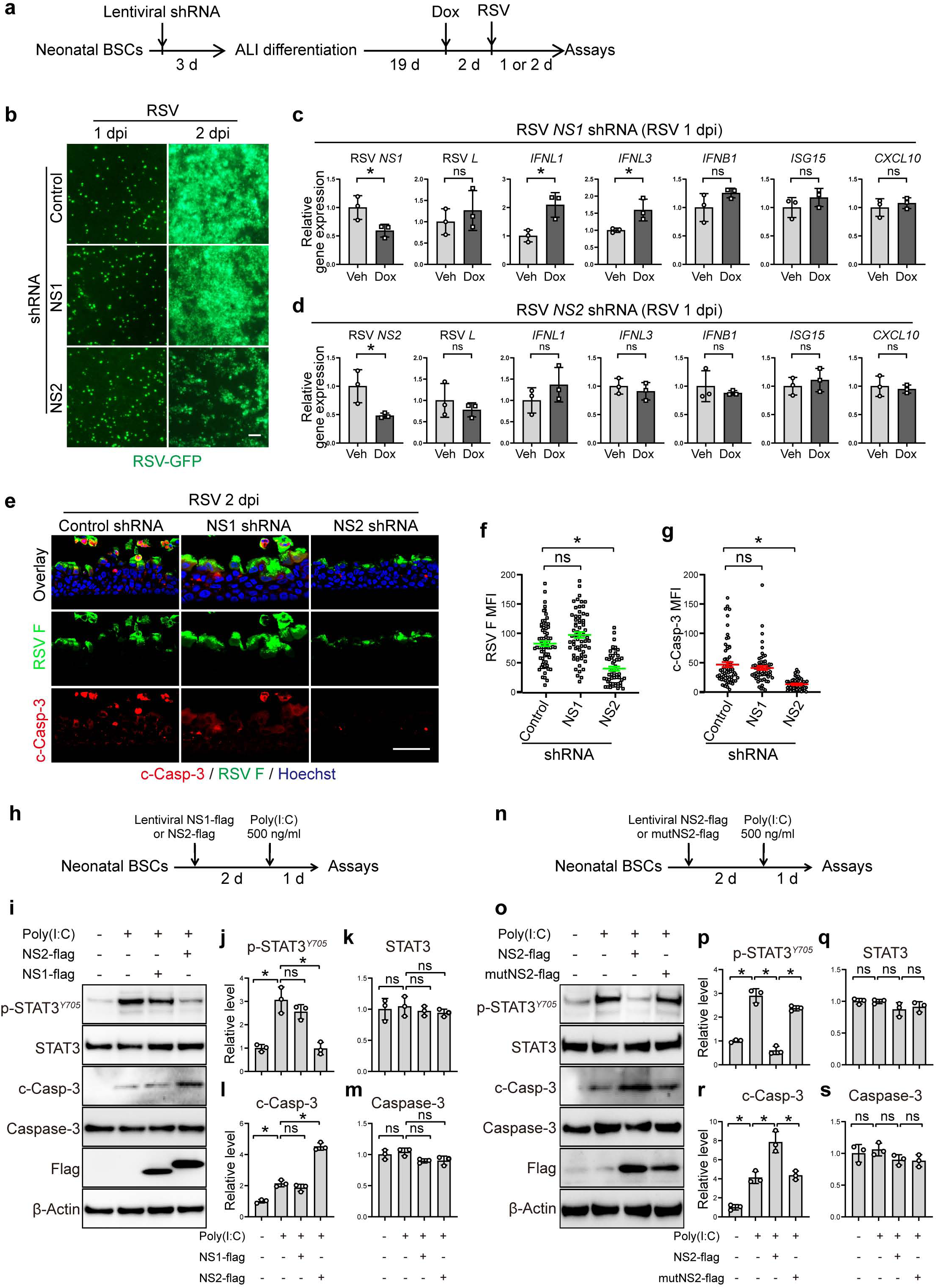
RSV NS2 is a critical viral mediator of severe RSV infection in neonatal BEpiCs. **a**, Schema of NS1 and NS2 knockdown by an inducible lentiviral shRNA system. Day 21 ALI cultures of lentivirus-transduced neonatal BSCs (N=3 donors) were treated with Dox (500 ng/mL) in the bottom chamber starting from day 19, infected with RSV A2-GFP (MOI 2) on day 21, and analyzed at 1 and 2 dpi. **b**, Representative images of GFP at 1 and 2 dpi. **c**,**d**, Relative mRNA levels of RSV *NS1, NS2*, and *L* genes and host *IFNL1*, *IFNL3, IFNB1*, *ISG15*, and *CXCL10* genes at 1 dpi by RT-qPCR. **e**, Representative double staining for RSV F and c-Casp-3 using sections of ALI cultures at 2 dpi. **f**,**g**, MFI of RSV F and c-Casp-3. Each dot represents one 20x image. At least 10 images from each donor were quantified. **h**, Schema of NS1-flag and NS2-flag overexpression in neonatal BSCs followed by treatment with poly (I:C) in liposome for 1 day and protein assays. **i**, Representative Western blot assays of p-STAT3^Y705^, STAT3, c-Casp-3, Caspase 3, and Flag tag in neonatal BSCs under each condition. β-Actin was loading control. **j-m**, Quantification of Western blot results by densitometry normalized to β-Actin. **n,** Schema of NS2-flag and mutant NS2 (mutNS2)-flag overexpression in neonatal BSCs followed by poly (I:C) treatment for 1 day and protein assays. **o**, Representative Western blot assays of p-STAT3^Y705^, STAT3, c-Casp-3, Caspase 3, and Flag tag in neonatal BSCs under each condition. **p-s**, Quantification of Western blot results by densitometry normalized to β-Actin. Bar graphs show mean ± SEM. Each mark represents one donor. *p<0.05 and ns, not significant by Student’s t-test (two-tailed) in **c**,**d** and one-way ANOVA followed Tukey’s multiple comparison test in **f**, **g**, **j**-**m**, and **p**-**s**. Scale bars, 50 μm.

Because BSCs are the cell origin of the immune-poised state in adult ciliated cells (Fig. 1) and can also be infected by RSV^42^, we employed neonatal and adult BSCs as a model to investigate how age regulates NS2-host interaction to determine the p-STAT3^Y705^ response to RSV. Neonatal BSCs were transduced to overexpress RSV NS2 carrying an N-terminus flag-tag to facilitate detection (Fig. 2h)^33^. RSV NS1, which is dispensable for severe infection of neonatal BEpiCs (Fig. 2b-g), was similarly overexpressed. Because overexpressed NS2 or NS1 may affect RSV replication and/or spread to alter host response, we simulated RSV infection using poly(I:C), a dsRNA analog. NS1 overexpression had no effect on neonatal BSC responses to poly(I:C), manifested by elevated levels of p-STAT3^Y705^ and cleaved Caspase 3 (c-Casp-3, a mark of apoptosis) (Fig. 2i,j,l). In contrast, NS2 overexpression diminished the p-STAT3^Y705^ response to poly(I:C) and exacerbated apoptosis. In addition, NS2 with 5 alanine (A) substitutions (24/25/26/32/33A) in the helix α1 domain (mutNS2)^33^ completely lost the ability to block the p-STAT3^Y705^ response to poly(I:C) and induce apoptosis (Fig. 2n-s). The levels of STAT3 and Caspase-3 were unchanged in any experimental condition (Fig. 2h-s). Therefore, RSV NS2 impairs the p-STAT3^Y705^ response to a viral analog by interacting with host factors via the helix α1 domain.

### RSV NS2 interacts with RIG-I in an age-associated manner to regulate RSV resistance in BEpiCs

The helix α1 domain of RSV NS2 is required for specific binding to RIG-I and MDA5 and subsequent inactivation^33^. Because only RIG-I shows age-associated expression in BEpiCs (Fig. 1) and is known to play an essential role in STAT3 activation in response to viruses^36^, we tested whether robust RIG-I levels in adults may outcompete RSV NS2 binding by co-immunoprecipitation (Co-IP) using the BSC model (Fig. 3a). RIG-I was barely pulled down in BSCs that overexpressed mutNS2 (Fig. 3b), providing direct evidence for an essential role of α1 helix in RSV NS2-RIG-I binding in human BSCs. Neonatal and adult BSCs, which overexpressed RSV NS2 at similar levels, showed no difference in RIG-I levels in protein pulldowns (Fig. 3c,d). However, RIG-I in the supernatants, which represented the pool of RIG-I not bound by NS2 and thus potentially available to activate the downstream signaling pathways, was more abundant in adult samples than neonatal samples (Fig. 3d,e). Therefore, elevated levels of RIG-I with age in adult BSCs can outcompete RSV NS2 binding.

**Fig. 3.**
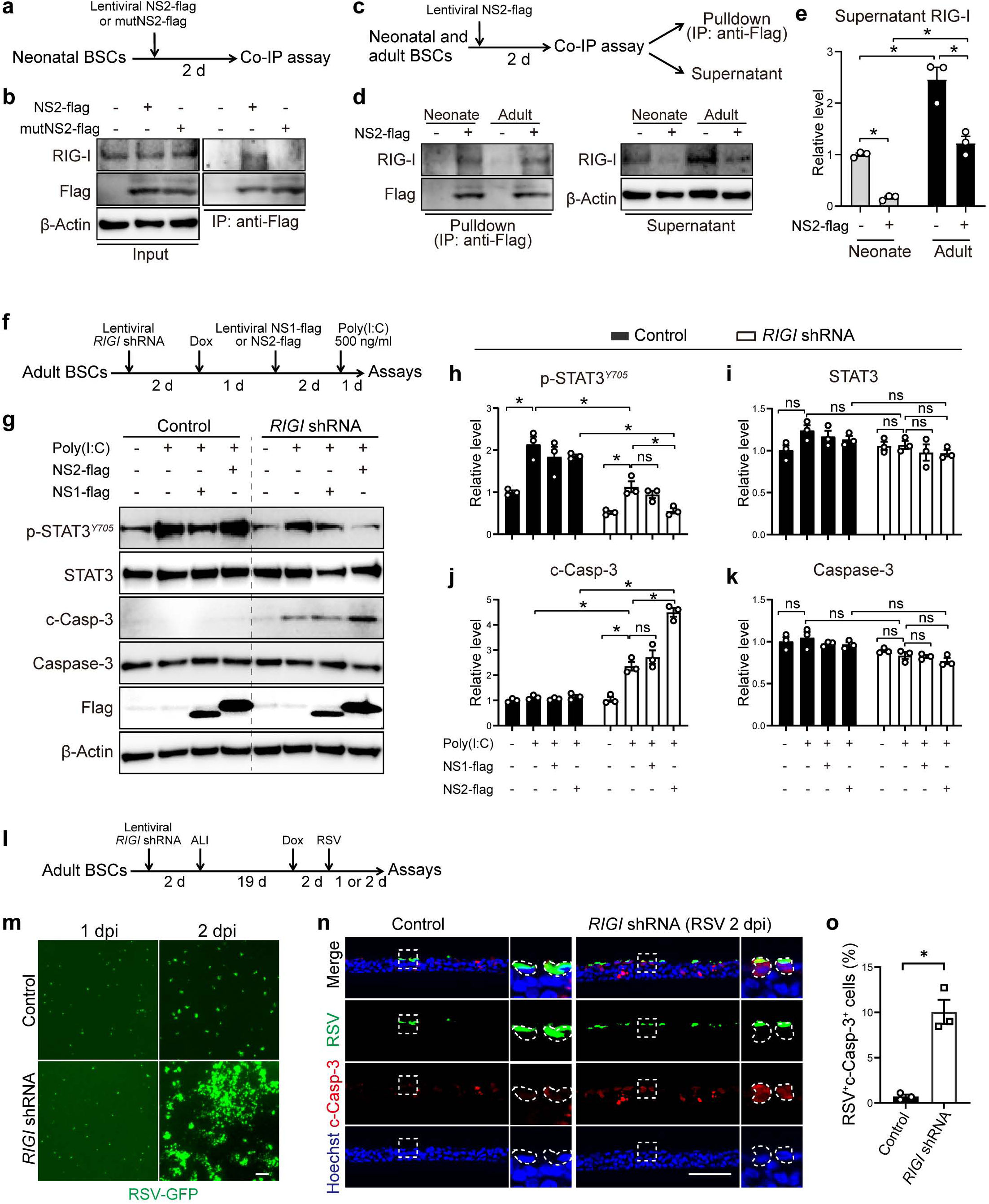
RSV NS2 binds to RIG-I to impair the p-STAT3^Y705^ response to poly(I:C) in an age-dependent manner. **a**, Schema of NS2-flag and mutant NS2(mutNS2)-flag overexpression in neonatal BSCs (N=3 donors) for 2 days followed by co-IP using the antibody against Flag tag. **b**, Representative Western blots of RIG-I and Flag tag in input and pulldown samples. **c**, Schema of NS2-flag overexpression in neonatal and adult BSCs followed by co-IP. **d**, Representative Western blots of pulldown and supernatant samples. **e**, Densitometry of Western blot results of the supernatant samples normalized to β-Actin. **f**, Schema of NS1 and NS2 overexpression in adult BSCs with Dox-inducible *RIGI* knockdown followed by poly (I:C) treatment for 1 day and protein assays. **g**, Representative Western blots for p-STAT3^Y705^, STAT3, c-Casp-3, Caspase 3, and Flag tag. β-Actin was loading control. **h-k**, Densitometry of Western blot results normalized to β-Actin. **l**, Schema of Dox-inducible *RIGI* knockdown in adult BSCs followed by differentiation in ALI, Dox (500 ng/mL) treatment from day 19, RSV A2-GFP infection on day 21, and assays at 1 and 2 dpi. **m**, Representative images of RSV-GFP-infected adult ALI cultures with and without *RIGI* knockdown. **n**, Representative images of double staining for RSV F and c-Casp-3 at 2 dpi. Scale bars, 50 μm. **o**, The relative abundance of RSV F^+^c-Casp-3^+^ cells in control and *RIGI* knockdown ALI cultures. Bar graphs show mean ± SEM. Each mark represents one donor. *p<0.05 and ns, not significant by two-way ANOVA followed by Sidak’s multiple comparison test in **e**,**h-k** and by Student’s t-test (two-tailed) in **o**.

To test whether the age-associated increase in RIG-I enables the p-STAT3^Y705^ response to RSV, adult BSCs were subjected to shRNA knockdown that reduced *RIGI* expression by ∼60% to a level comparable to that in neonatal BSCs (Fig. 3f and Extended Data Fig. 3). Control adult BSCs transduced with scrambled shRNA showed a robust p-STAT3^Y705^ response to poly(I:C) and undetectable levels of c-Casp-3 regardless of NS1 or NS2 overexpression (Fig. 3g,h,j). However, *RIGI* knockdown dampened the p-STAT3^Y705^ response to poly(I:C) and increased apoptosis, and these changes by *RIGI* knockdown were worsened by overexpressed NS2, but not NS1. *RIGI* knockdown had no effect on STAT3 and Caspase 3 (Fig. 3g,I,k). These findings show that adult BSCs with *RIGI* knockdown resemble neonatal BSCs in response to RSV infection simulated by poly(I:C) with NS2 overexpression.

To test whether age-associated RIG-I upregulation renders RSV resistance in adult BEpiCs, adult BSCs with *RIGI* knockdown differentiated in ALI for 3 weeks followed by infection with RSV-GFP (MOI 2) (Fig. 3i). RIG-I knockdown had no effect on RSV infection or the host IFN response at 1 dpi (Fig. 3m and Extended Data Fig. 3), consistent with a marginal role of RIG-I in the anti-viral response in BEpiCs (lower airway)^35^. However, at 2 dpi, RIG-I knockdown in adult BEpiCs significantly promoted virus spread and apoptosis (Fig. 3m-o). Treatment of adult ALI cultures with RIG012, a small molecule RIG-I inhibitor, also worsened RSV infection without affecting the host IFN response (Extended Data Fig. 3). Therefore, RIG-I, which is elevated with age, plays a critical role in RSV resistance in adult BEpiCs.

### Baseline p-STAT3^Y705^ partners with increased chromatin accessibility to sustain RIG-I upregulation with age

The age-associated immune-poised state in adult BEpiCs in the human lung was sustained *in vitro (*Fig. 1), suggesting epigenetic regulation. To test this hypothesis, we performed ATAC-seq of neonatal and adult BSCs and detected age-associated chromatin accessibility around the transcription start site (TSS) in adult BSCs (Fig. 4a). Consistently, Western blot and antibody staining showed a reduction in a repressive histone mark, H3K27me3, in adult BSCs and BEpiCs compared to the infant counterparts (Fig. 4b-f). Age had no effect on two activating histone marks, H3K27ac and H3K4me3 (Fig. 4b-e,g). Therefore, age is associated with increased chromatin accessibility and a decrease in H3K27me3 in adult BSCs and progeny BEpiCs.

**Fig. 4.**
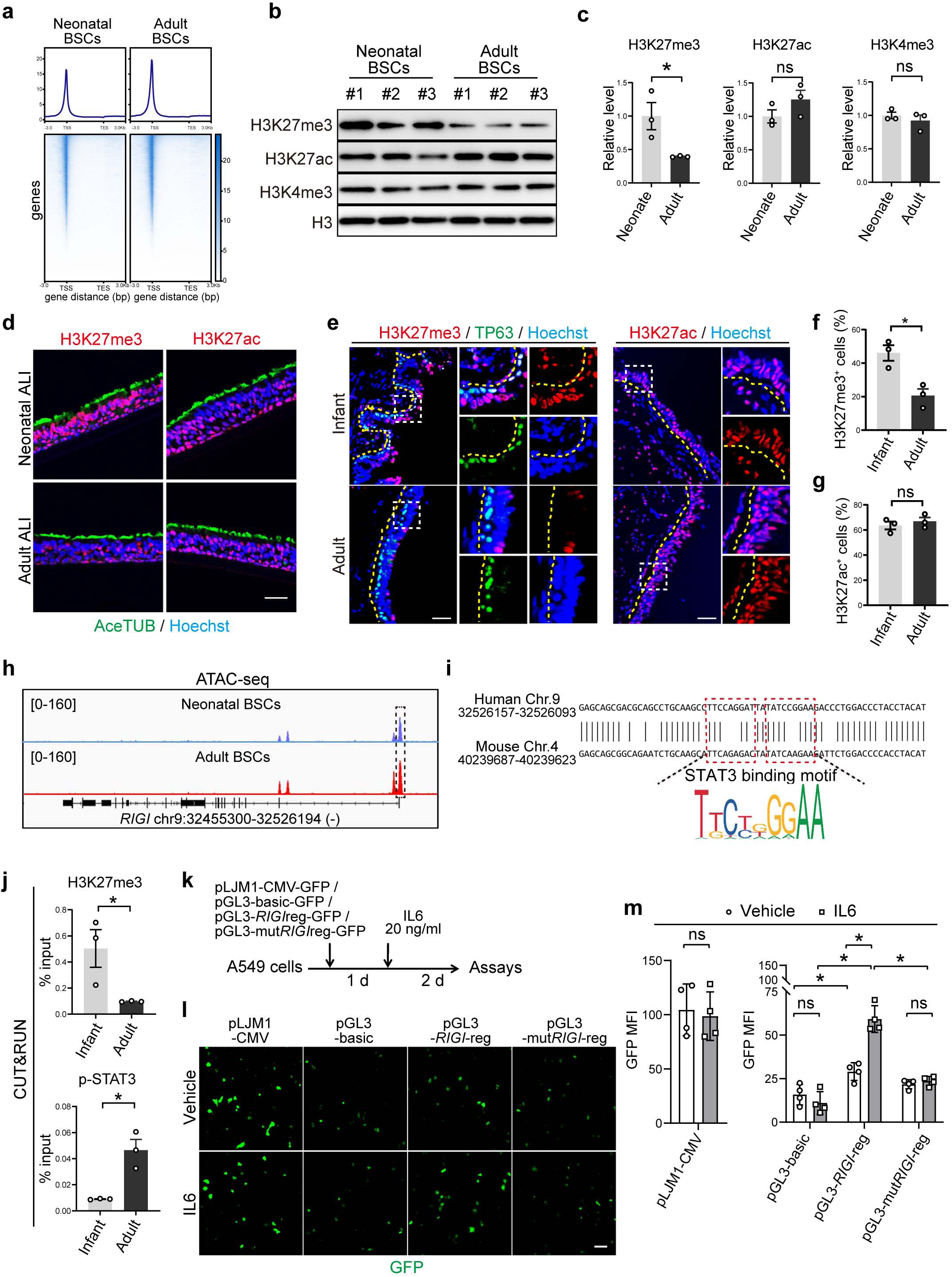
Age-associated upregulation of RIG-I in adult BEpiCs is sustained by baseline p-STAT3^Y705^ and increased chromatin accessibility. **a**, Heatmap of ATAC-seq peaks around the transcription start site (TSS) in neonatal and adult BSCs. **b**, Representative Western blots of histone marks in neonatal and adult BSCs. H3 was loading control. Each lane is one donor. **c**, Densitometry of Western blot results normalized to H3. **d**,**e**, Representative images of staining for H3K27me3 and H3K27ac in BEpiCs in ALI cultures (e) and donor lungs of infants and adults (N=3 per age group). Some images showed co-staining for AceTUB (acetylated tubulin, a ciliated cell marker) and TP63 (a BSC marker). Nuclei were counter stained with Hoechst dye. The yellow dotted line marks basement membrane of the bronchial epithelium. **f**,**g**, The relative abundance of H3K27me3^+^ and H3K27ac^+^ cells in the bronchial epithelium in infant and adult donor lungs. Each mark is one donor. **h**, Peaks of *RIGI* gene accessibility in neonatal and adult BSCs. **i**, Conserved regulatory sequences (*RIGI*reg) within the outlined peak in **h**. **j**, H3K27me3 and p-STAT3^Y705^ binding to *RIGI*reg by CUT&RUN in neonatal and adult BSCs (N=3 per age group). **k**, Schema of transcriptional activity assays of *RIGI*reg in A549 cells using a pGL3-GFP reporter with IL6 treatment (20 ng/mL). Point mutations to disrupt the STAT3 binding motif were introduced (mut*RIGI*reg). **l**, Representative images of GFP after 48 hours of IL6 treatment. **m**, MFI of GFP in 4 independent experiments. Bar graphs show mean ± SEM. *p<0.05 and ns, not significant by Student’s t-test (two-tailed) in **c**,**f**,**g**,**j** and two-way ANOVA followed by Sidak’s multiple comparison test in **m**. Scale bars, 50 μm.

We detected four ATAC-seq peaks near the 5’ end and within the introns of the human *RIGI* gene, all of which showed increased intensity with age (Fig. 4h). However, only the peak at the 5’ end (chr9: 32,526,907-32,526,346 (-)) contains evolutionally conserved DNA sequences that overlap with a known active enhancer^43^ containing two tandem repeats of the STAT3 binding motif (Fig. 4i). CUT&RUN with a H3K27me3 antibody following by qPCR using primers flanking the conserved STAT3 binding motif showed reduced H3K27me3 and reciprocally increased p-STAT3^Y705^ binding to the conserved regulatory sequences (*RIGI*reg) in adult BSCs compared to neonatal BSCs (Fig. 4j). Because adult BEpiCs have elevated baseline p-STAT3^Y705^ compared to neonatal BEpiCs (Fig. 1c-f,h)^24^, increased accessibility to *RIGI*reg with age may facilitate *RIGI* gene expression driven by p-STAT3^Y705^.

To test this hypothesis, we first assessed the effect of STAT3 blockade on RIG-I levels in adult BEpiC using a small molecule inhibitor Stattic and a dominant negative STAT3-S560del mutation in Job syndrome. Both approaches abolished RSV resistance in adult BEpiCs^24^ and reduced the level of RIG-I (Extended Data Fig. 4). In addition, the STAT3-S560del mutation reversed age-associated enrichments in the inflammatory response pathways in adult BEpiCs without affecting the reduction in H3K27me3 with age (Extended Data Fig. 4), suggesting that epigenetic changes with age in human BEpiCs is independent of active STAT3. Furthermore, we evaluated the transcriptional activity of *RIGI*reg by inserting the sequences into a basic GFP reporter vector (pGL3-basic-GFP). After transfection into an A549 lung cancer cell line carrying active STAT3^44,45^, *RIGI*reg significantly elevated GFP reporter expression, compared to the basic vector, at baseline and in response to IL6 treatment (Fig. 4k-m). However, these activities were completely abolished by point mutations that disrupted the two STAT3 binding motifs (mut*RIGI*reg). As control, we showed that IL6 treatment had no effect on GFP expression from the pGL3-basic vector or driven by a constitutively active cytomegalovirus (CMV) promoter (Fig. 4k-m). Therefore, age-associated baseline p-STAT3^Y705^ partnering with chromatin accessibility plays an essential role in driving the sustained immune-poised state, including elevated RIG-I levels, in adult BEpiCs.

### Acute respiratory viral infection induces a persistent immune-poised state in mouse BEpiCs

To test whether the immune-poised state of adult BEpiCs in humans reflects the outcome of postnatal development associated with physical age, we re-analyzed available scRNA-seq datasets that compared lungs from neonatal, adult (3-month), and aged (24-month) mice housed in specific pathogen-free (SPF) facility^46–49^. Physical age was not associated with the enrichments of the inflammatory response pathways in BEpiCs in mice (Extended Data Fig. 5).

Different from SPF mouse lungs, human lungs are constantly challenged by pathogens. We thus tested whether acute respiratory viral infection in mice can induce a similar immune-poised state in BEpiCs found in humans with age. In a published scRNA-seq dataset of a mouse influenza A virus (IAV) infection model (Fig. 5a)^46^, a total of 70,770 lung cells from four experimental groups (mock, 6 dpi, 42 dpi, and 366 dpi) were clustered and each group included over 10,000 single cells from 2-4 mice (Extended Data Fig. 6). At the peak of infection around 6 dpi in mice^46,50^, robust induction of genes involved in active infection and inflammation dominated the list of the 15 most significantly enriched pathways in BEpiCs (Fig. 5b and Extended Data Fig. 6). At 42 dpi when IAV is cleared and lung repair is near completion, changes in genes involved in the IFN production, viral replication, and antiviral innate immune response pathways were resolved. However, the enrichments in the IFN, IL6, and TNFα response pathways in BEpiCs remained significant at 42 dpi and even after 1 year when lung homeostasis has been re-established (Fig. 5b). In contrast, AT1 and AT2 cells showed a transient inflammatory gene signature only at 6 dpi. These findings indicate that respiratory viral infection in mice exerts a long-term impact on gene expression in an epithelial cell type-specific manner, similar to our findings in human lungs with age (Fig. 1).

**Fig. 5.**
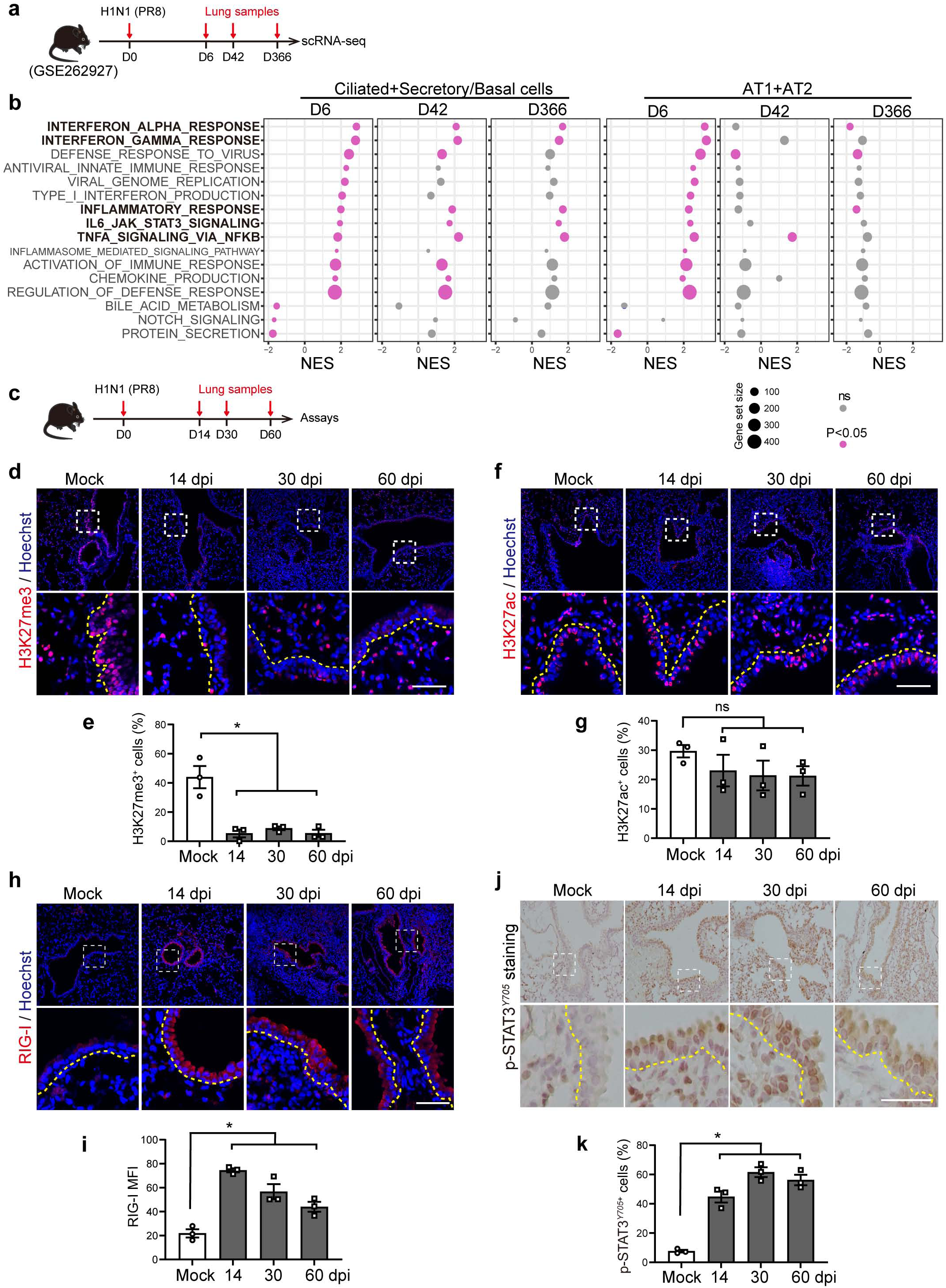
Acute respiratory viral infection in SPF mice induces immune-poised characteristics in BEpiCs. **a**, Schema of mouse flu infection and scRNA-seq in a public dataset (GSE262927). **b**, Dot plots of enriched hallmark pathways in BEpiC and AT1+AT2 cells at each time point after infection. Mock group is baseline control. **c**, Schema of mouse flu infection for lung section staining. **d**,**f**,**h**,**J**, Representative images of fluorescence staining for H3K27me3, H3K27ac and RIG-I and chromogenic staining for p-STAT3^Y705^ in mock and H1N1-infected mouse lungs. Cells residing in the lung mesenchyme within the same images, which showed comparable staining between mock and IAV infection, were control for BEpiC-specific changes following infection. The bottom panels show enlarged outlined areas in the upper panels. The dotted line marks basement membrane. Nuclei were stained by Hoechst dye in fluorescence staining and by hematoxylin in chromogenic staining. Scale bars, 50 µm. **e**,**g**,**i**,**k**, Quantification by MFI or the percentage of positive epithelial cells. Bar graphs show mean ± SEM. Each dot represents one mouse. *p<0.05 and ns, not significant by one-way ANOVA followed Tukey’s multiple comparison tests.

We showed no change in robust H3K27me3 in BEpiCs in neonatal and adult mice (Extended Data Fig. 7). Following IAV infection in adult mice^51^, the relative abundance of H3K27me3^+^ BEpiCs was reduced as early as 14 dpi and remained at a low level at least until 60 dpi (Fig. 5c-e). In contrast, IAV infection had no effect on H3k27ac in BEpiCs (Fig. 5f,g). In addition, the level of RIG-I and the relative abundance of p-STAT3^Y705^–positive cells in the mouse bronchial epithelium were increased at multiple time points after IAV infection (Fig. 5h-k). Cells in the lung mesenchyme had comparable levels of H3K27me3, RIG-I, and p-STAT3^Y705^ regardless of IAV infection, providing further evidence for BEpiC-specific influence of IAV infection. Similar differential changes in H3K27 marks, RIG-I, and p-STAT3^Y705^ in mouse BEpiCs were found following SARS-CoV-2 infection^52^ (Extended Data Fig. 7). Taken together, acute infection by different respiratory viruses induces sustained changes in mouse BEpiCs that resemble key features of the immune-poised physiological state in human BEpiCs with age.

### Cytokine pretreatment of neonatal BSCs induces adult-like RSV resistance in progeny BEpiCs

To test whether inflammation triggered by respiratory viral infection during a lifetime conveys the effect of age on BSCs and progeny BEpiCs, we treated neonatal BSCs (never exposed to inflammation) with cytokines at a physiologically relevant concentration of 2 ng/mL for 7 days, an average duration of acute respiratory viral infection (Fig. 6a and Extended Data Fig. 8). At this low concentration, individual cytokine or cytokine cocktail (IL6+TNFa+IFNγ) tested had a marginal effect on BSC proliferation. We reasoned that epigenetic changes found in adult BEpiCs with age may occur as an early event to sustain the inflammatory response gene signature. Therefore, we assessed changes in H3K27 modification in BSCs immediately after 7-day treatment as an early indicator of efficacy. Compared to single cytokine treatment that had no effect, cytokine cocktail treatment significantly reduced the level of H3K27me3 in neonatal BSCs (Extended Data Fig. 8) and the reduction was maintained over two passages (each at a 1:4 split) in the absence of continuous treatment (Fig. 6b,c). Neonatal BSCs with cytokine cocktail pretreatment also showed increased levels of RIG-I and p-STAT3^Y705^ that were associated with reduced H3K27me3 and increased p-STAT3^Y705^ binding to the *RIGI*reg sequences (Fig. 6b-d). No change in H3K27ac or STAT3 was found under any treatment conditions (Fig. 6b,c and Extended Data Fig. 8). Furthermore, we found no difference in differential potentials of cocktail-pretreated neonatal BSCs compared to vehicle-pretreated control in ALI (Fig. 6e,f). Following RSV-GFP infection, ALI cultures generated from cytokine cocktail-pretreated neonatal BSCs had a more than 4-fold decrease in the abundance of RSV^+^ apoptotic ciliated cells at 2 dpi compared to vehicle-pretreated control (Fig. 6f,g). Lastly, to link the NS2-RIG-I-STAT3 axis to RSV resistance induced by cytokine cocktail pretreatment in neonatal BEpiCs, we employed the Dox-inducible shRNA approach to knockdown RIG-I. Compared to scrambled shRNA control, RIG-I knockdown in ALI cultures almost completely abolished RSV resistance induced by cytokine cocktail pretreatment (Fig. 6e-g). In addition, vehicle-treated neonatal BSCs showed increases in p-STAT3^Y705^ and c-Casp-3 in response to poly(I:C) and RSV NS2 dampened the p-STAT3^Y705^ response and elevated apoptosis (Fig. 6h-m). In comparison, cytokine cocktail-pretreated neonatal BSCs resembled adult BSCs in response to poly(I:C), evidenced by a robust increase in p-STAT3^Y705^ above a higher baseline and resistance to apoptosis even in the presence of RSV NS2 overexpression (Figs 6h-m). In contrast, RSV NS1 overexpression had no effect on neonatal BSC responses to poly(I:C) regardless of cytokine cocktail pretreatment. These findings indicate that cytokine cocktail pretreatment of neonatal BSCs induces an adult-like immune-poised state and renders RSV resistance in differentiated progeny cells through the RSV NS2-RIG-I-STAT3 axis.

**Fig. 6.**
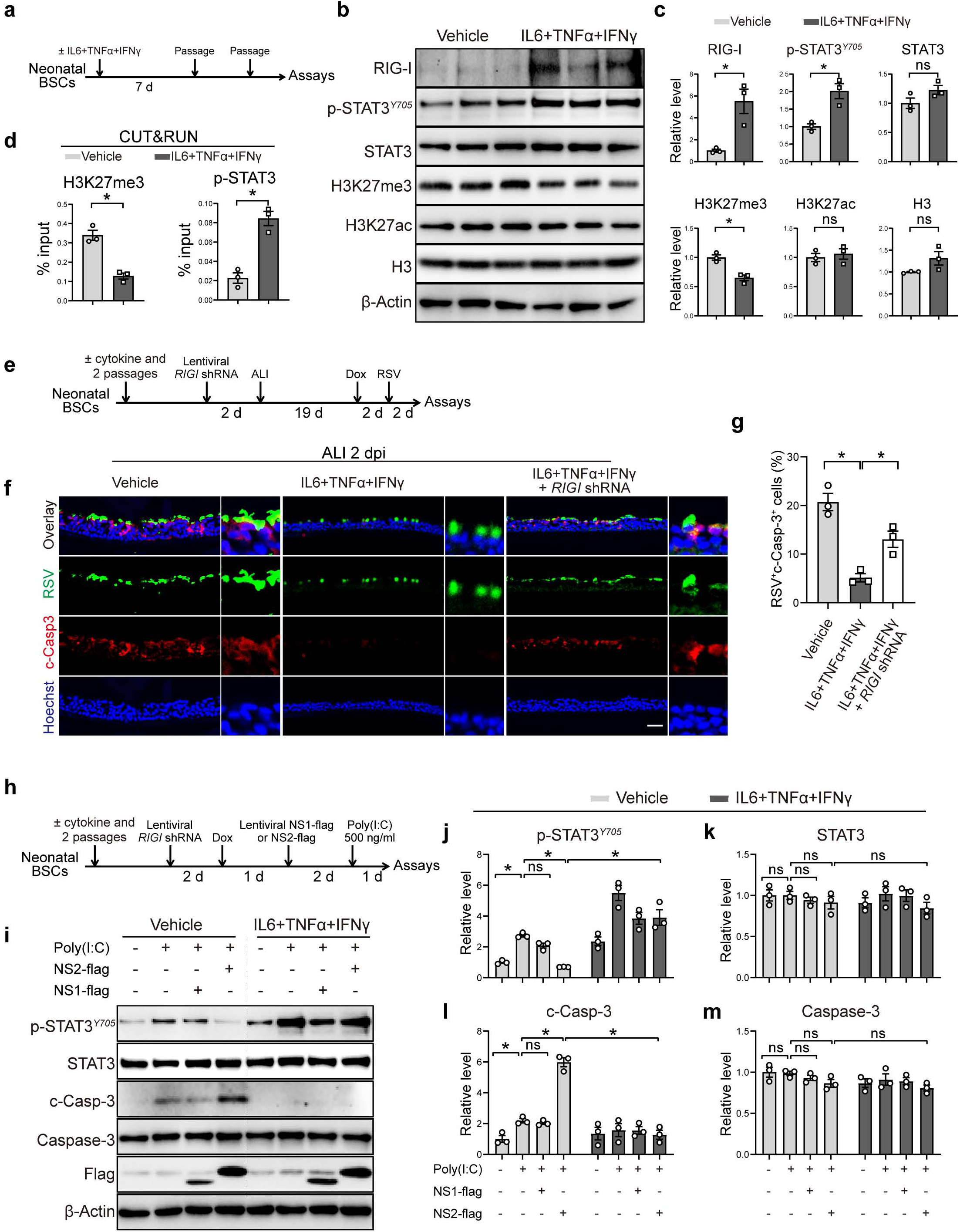
Cytokine cocktail pretreatment of neonatal BSCS induces adult-like RSV resistance in progeny BEpiCs. **a**, Schema of 7-day cytokine cocktail (IL6+TNFα+IFNγ, each at 2 ng/mL) treatment of neonatal BSCs followed by two passages (each at a 1:4 split). **b**, Representative Western blots of RIG-I, p-STAT3^Y705^, and H3K27 modifications in neonatal BSCs pretreated with vehicle or cytokine cocktail. β-Actin was loading control. Each lane is one donor. **c**, Densitometry of Western blot results normalized to β-Actin. **d**, H3K27me3 and p-STAT3^Y705^ binding to the regulatory sequences of the *RIGI* gene in vehicle-and cytokine cocktail-pretreated BSCs by CUT&RUN. **e**, Schema of Dox-inducible *RIGI* knockdown in vehicle- or cytokine cocktail-pretreated neonatal BSCs followed by differentiation and RSV A2-GFP infection. **f**, Representative images of double staining for RSV F and c-Casp-3 in ALI cultures at 2 dpi. Scale bar, 50 µm. **g**, Relative abundance of RSV F^+^c-Casp-3^+^ cells quantified from double stained images. Each mark is one donor. **h**, Schema of NS1 and NS2 overexpression, *RIGI* knockdown and poly(I:C) treatment of neonatal BSCs pretreated with vehicle or cytokine cocktail. **i**, Representative Western blot assays for p-STAT3^Y705^, STAT3, c-Casp-3, Caspase 3, and Flag tag in each condition. β-Actin was loading control. **j**-**m**, Densitometry of Western blot results normalized to β-Actin. Each mark represents one donor. Bar graphs show mean ± SEM. *p<0.05 and ns, not significant by Student’s t-test (two-tailed) in **c**,**d**, one-way ANOVA followed Tukey’s multiple comparison test in **g**, and two-way ANOVA followed by Sidak’s multiple comparison test in **j**-**m**.

## DISCUSSION

Building upon our previous findings that a STAT3 activation response to RSV in the human bronchial epithelial barrier critically determines the severity of RSV infection with age^24^, this study determines RSV NS2 interaction with host factor RIG-I as a critical upstream regulator of the STAT3 response to RSV. In addition, we show that respiratory infection during a lifetime can prime RSV resistance by inducing an immune-poised physical state in human BSCs followed by transmission to differentiated progeny cells, including ciliated cells, which uncovers a mechanism of age in regulating RSV susceptibility in humans. Furthermore, we find that the immune-poised state is sustained by a transcriptional program driven by baseline p-STAT3^Y705^ and epigenetic reprogramming, which fortifies RIG-I to guard against RSV NS2-mediated binding and inactivation. This role of RIG-I in age-related RSV susceptibility is independent of the classical dsRNA sensor activity of RIG-I in triggering the anti-viral IFN response following infection. Instead, RIG-I triggers Syk-mediated STAT3 phosphorylation in response to viral infection^36^. Lastly, by demonstrating that the unique α1 helix of RSV NS2 is specifically required for binding to RIG-I and blocking the RIG-I-STAT3 pathway, our findings provide an explanation for RSV specificity in age-related resistance in the human bronchial epithelial barrier. Notably, NS2 knockdown in neonatal ALI cultures only partially restored RSV resistance found in adults^24^, suggesting that the remaining RSV NS2 and/or other viral factors contribute to age-related RSV-host ciliated cell interaction to exacerbate the severity of infection.

How age differentially regulates the level of baseline p-STAT3^Y705^ and H3K27me3 in human BEpiCs warrants future investigation. Based on our findings of respiratory infection in specific pathogen-free mice and inflammatory cytokine treatment of human neonatal BSCs, infection and inflammation during a lifetime may mediate the effect of age on human BEpiCs. In support of this hypothesis, BSCs directly communicate with resident innate immune cells and resident memory T cells in the lung^53,54^. The link between acute respiratory infection and inflammation and the immune-poised state in BEpiCs aligns with the concept of inflammatory memory found in tissue-specific stem cells^55^. In multiple organs, including the skin, the gut, and the pancreas, transient injury and/or inflammation cause sustained STAT3 activation and chromatin accessibility in tissue-resident stem cells to induce memory, and such memory serves to boost the stem cell function in barrier restoration following a subsequent insult^56–58^. Mechanisms of memory induction in these tissue-specific stem cells are similar to the effect of age on human BEpiCs found in this study. These previous studies also show that apart from priming stem cells in tissue repair, inflammatory memory in tissue-specific stem cells increases susceptibility to cancer and chronic inflammatory diseases. Whether the immune-poised physiological state predisposes adult BEpiCs to pathogenic changes in lung cancer or chronic airway diseases is unknown. However, we recently reported that excessive cytokine treatment of healthy adult BSCs leads to mucus hyperplasia after differentiation^23^, suggesting that the immune-posed physiological state in adult BSCs may be exploited by subsequent severe respiratory viral infection to contribute to long term sequelae of infection.

Due to a lack of *in vivo* models of severe RSV infection, whether an early life acute infection in the lung protects against severe RSV infection in adulthood has yet to be fully assessed. A recent retrospective cohort study of children aged 0-5 years reported that prior COVID-19 infection increases risk for RSV infections^59^. Although the conclusion from this clinical study may appear to contradict our conclusion of infection-induced RSV protection, we need to point out that the incidence of other common viral infections in the study groups was unreported, which may contribute to the low RSV infection rate in the non-COVID group.

In summary, our findings inform fundamental biology of age in the bronchial epithelial barrier that underlies RSV susceptibility in humans. The bolstered barrier protection in adults likely orchestrates and partakes in the action of mature immune cells to protect against severe RSV infection. The precise age of transition from vulnerability in young children to resistance in adults and whether and how genetic and environmental factors affect the age of transition warrant future investigation and may yield valuable markers to identify high-risk individuals for additional protection. Strategies to induce an adult-like RSV NS2-RIG-I-STAT3 axis following RSV infection may provide effective treatment that complement clinically available RSV vaccines and antibodies and in combination, offer full protection against RSV in infants.

## Supporting information

All supplemental files

## ACKNOWLEDGEMENTS

This work was supported by NIH grants to XA (R21AI173494) and DWL (R01AI159678) and funds from the Department of Pediatrics at MGH for Lung Cell Bank to XA. We would like to thank all members of the Ai lab and Dr. Bryan P. Hurley at MGH for constructive criticisms.

## AUTHOR CONTRIBUTIONS

C.Z. performed RSV infection experiments and analyzed the transcriptomic data. P.H.L. and H.M. provided BSCs from human patients. X.W., H.N., and J.S. provided lung sections of mouse infection models. D.W.L. provided DNA vectors for RSV NS1 and NS2. J.X. provided methodological guidance. R.F. guided RSV infection experiments. X.A. and C.Z. conceived the study. C.Z. and X.A. wrote the manuscript. Y.B. edited the manuscript. All the authors read and commented on the manuscript.

## Methods

### BSC isolation and expansion

BSC derivation from TA samples of selected patients was described previously^20,21^. Briefly, TA samples were collected from intubated neonatal and adult patients at the Intensive Care Unit (ICU), Massachusetts General Hospital (MGH) under an approved IRB protocol (No. 2019P003296). Neonatal and adult patients for TA sample collection were intubated due to cardiogenic and neurogenic respiratory failure with no known lung diseases. BSCs in TA samples were cultured and expanded in Small Airway Epithelial Cell Growth Medium (SAGM, PromoCell, Cat# C-21070) in the presence of SMAD/ROCK/mTOR inhibitors^20–22^. BSCs of one patient with Job syndrome were isolated from fresh discarded surgical specimens under an approved IRB protocol (No. 2017P001479) as previously reported^24,60^ and expanded under similar culture condition as TA BSCs.

### RSV preparation

RSV A2 (VR-1540) was purchased from ATCC (VR-1400). RSV A2-GFP strain was purchased from ViraTree (https://www.viratree.com/). RSV was propagated in Hep-2 cell line (ATCC CCL-23). Hep-2 cells were cultured in Opti-MEM (Gibco) supplemented with 2% fetal bovine serum (FBS) (Gibco) and 1% GlutaMAX Supplement (Gibco). Hep-2 cells at 80% confluency were added and infected with RSV in complete medium at a multiplicity of infection (MOI) of 0.1 for 1 h at 37°C. When high cytopathic effects were observed (∼5 days post infection) in culture, cells were scraped into the medium, cell debris was separated by centrifugation, and the supernatant was aliquoted and stored at −80°C. RSV virus titers were measured via plaque assay as previously described^61^.

### Mucociliary differentiation of BSCs in ALI

As previously described^20^, 2×10^5 BSCs were seeded in 24-well 6.5 mm Transwell inserts with 0.4 μm polyester membranes (Corning, Cat#3470), which were precoated with stored 804G-conditioned medium for overnight. Differentiation in PneumaCult medium (StemCell Technologies, Cat#05001) was initiated by removing apical medium and maintained at air-liquid interface (ALI) for 21 days, with medium changes every 2 days. On day 21, ALI cultures were used for viral infection assays, followed with fixation for downstream antibody staining or assays. For Dox-inducible NS1/NS2/RIGI knockdown experiments, BSCs were first selected with puromycin (1 μg/mL) for 3 days before ALI differentiation. On day 18 of ALI, cultures were treated with vehicle or Dox (500 ng/mL) in the basal medium for 3 days, and knockdown efficiency and RSV infection assays were performed on day 21.

### Infection of ALI cultures with RSV

Frozen RSV A2-GFP aliquots were thawed and diluted in PBS to the desired titer immediately before use. Prior to infection, the apical surface of ALI cultures was washed twice with 200 μL PBS to remove secreted mucus. RSV inoculum (200 μL) was applied apically for 1 h at 37 °C, 5% CO₂ in incubator, followed by three warm PBS washes to remove unbound RSV particles. Cytopathic effects were monitored daily by bright-field microscopy or GFP observation. RSV-infected epithelial cells were either fixed in 4% paraformaldehyde/PBS for antibody staining or lysed for protein and RNA assays.

### Antibody staining of ALI cultures

Mock or RSV-infected ALI cultures were fixed for 15 min (RSV) in 4% paraformaldehyde/PBS before they were processed for cryosectioning at a thickness of 10 μm or whole-mount staining. Sections or whole-mount ALI cultures were immune-stained according to a standard protocol using primary antibodies and secondary antibodies listed in the resource table (Supplementary Data 1). Nuclei were stained with Hoechst dye (1:500, ThermoFisher, Cat# H3570). Secondary antibodies for immunofluorescent staining include goat anti-rabbit Alexa Fluor 594 (IgG, 1:200, Cat#A-11037, Thermo Scientific) and goat anti-mouse Alexa Fluor 488 (IgG, 1:200, Cat#A-11029, Thermo Scientific).

### Antibody staining of human and mouse lung tissue sections

De-identified donor lungs were obtained from the International Institute for the Advancement of Medicine (IIAM) were used for preparation of paraffin-embedded lung section. This study included the analysis of three adult lungs (one female, two males; ages 43–59) and three infant lungs (one female, two males; ages 0–6 months). All donors had no known history of lung disease. Because no interventions or interactions with living individuals were involved in this study, the MGH Institutional Review Board classified this project as non-human subject research. IAV infection and SARS-CoV-2 infection of mouse model were carried as previously reported^51,52^. All mice were maintained in a specific-pathogen-free facility and used in accordance with protocols reviewed and approved by the Institutional Animal Care and Use Committees of the University of Virginia (Charlottesville, VA). For IAV infection, mice (8-12 weeks) were intranasally inoculated with 75 plaque-forming units (PFU) of the influenza A/PR8/34 strain, which had been diluted in fetal bovine serum (FBS)-free Dulbecco’s modified Eagle’s medium (DMEM; Corning) on ice. For SARS-CoV-2 infection, male C57BL/6mice aged 5 to 6 months (male mice of this age range are more susceptible to infection) were intranasally inoculated with 2000 PFU of mouse-adapted strain SARS-CoV-2 MA30. IAV- and SARS-CoV-2 infected mice were sacrificed at indicated time point. The harvested lungs were perfused with ice-cold PBS, and then inflated and fixed in 10% formalin, embedded in paraffin, and sectioned at 5 µm thickness. Paraffin-embedded lung sections were then processed for immunostaining using standard protocols in the lab. Primary antibodies and secondary antibodies are listed in the resource table (Supplementary Data 1).

### Transcriptome Analysis of BSCs by bulk RNA-seq

Bulk RNA-seq service was provided by GENEWIZ (South Plainfield, NJ, USA) or Active Motif (Carlsbad, CA, USA). Total RNA (2 μg per sample) was extracted from 1×10^6 cultured TA BSCs using the RNeasy Kit (Qiagen, Cat#74104), and libraries were prepared with the TruSeq V2 Kit (Illumina, Cat#20020594) by company. Libraries were sequenced on an Illumina NextSeq 500 platform to generate paired-end 42-nt reads, which were aligned to the human hg38 reference genome using STAR (v2.6.0a). Differential expression and principal component analyses were performed with DESeq2 (v1.30.0) in R, and genes with p-values <0.05 were considered differentially expressed.

### CUT&RUN

CUT&RUN was carried out following manufacturer’s instructions (Cell Signaling Technology, Cat# 86652). Cultured BSCs (10^5) were fixed with 0.1% formaldehyde for 2 min and bound to Concanavalin A–coated magnetic beads, permeabilized with digitonin-containing wash buffer, and incubated overnight at 4 °C with primary antibody. After washes, Protein A–MNase was added for 1 h at 4 °C. Digestion was activated with CaCl_2_, then stopped with provided stop buffer. Released fragments were incubated for 10 min at 37 °C, added 3 µl of 10% SDS Solution (0.1% final concentration) and 2 µl of proteinase K (20 mg/mL) overnight at 65°C to reverse the crosslinks. DNA was purified using the DNA Purification Buffers and Spin Columns (Cell Signaling Technology, Cat# 14209). DNA was quantified by RT-qPCR with primers designed in specific chromatin opening region of *RIGI* gene. The CUT&RUN assay primer sequences were summarized in the primer table (Supplementary Data 2).

### ATAC-seq

ATAC-seq was performed by ActiveMotif (Carlsbad, CA, USA). BSCs (1×10^5) were tagmented using the Nextera Library Prep Kit (Illumina), and libraries were purified, PCR-amplified, and sequenced on an Illumina NextSeq 500 as paired-end 42-nt reads by company. Reads were mapped to the human hg38 genome using BWA (v0.7.12), filtered for quality and uniqueness, and duplicates removed. Alignments were extended to 200 bp and stored in bigWig format. Peaks were called with MACS2 (v2.1.0, p<1e-7, – nomodel) with ENCODE blacklist regions excluded from analysis. Read counts in peaks were quantified with Rsubread (v2.6.4), and differential accessibility was analyzed with DESeq2 (v1.30.0). Motif enrichment was performed using HOMER (v4.11), and ATAC-seq tracks were visualized with IGV_2.16.0 software through uploading bigWig files.

### Lentiviral packaging and BSC infection

For knockdown experiments, short hairpin RNA (shRNA) sequences targeting *NS1*/*NS2*/*RIGI* genes were cloned into the pLKO.1 plasmid. For overexpression experiments, codon-optimized NS1 and NS2 sequences added with a Flag sequence at the C-terminus were cloned into pLJM1 plasmid (Addgene, Cat# 19319). The mutated NS2 was constructed with Q5® Site-Directed Mutagenesis Kit (NEB, Cat# E0554S). The constructed vector together with packaging plasmids were co-transfected into 293T cells using Lipofectamine 3000 (Thermo Scientific, Cat#L3000015) and lentiviral supernatant was collected 48-and 72-h post-transfection, filtered, and centrifugated. BSCs (1×10^5, ∼90% confluency) were infected with 300 μL of lentiviral supernatant in the presence of 10 μg/mL polybrene (Merck Millipore, Cat# TR-1003-G) for 12 h, followed by treatment with 10 ng/mL puromycin. The knockdown assay primer sequences targeting *NS1*/*NS2*/*RIGI* genes were summarized in Supplementary Table 1.

### Co-immunoprecipitation (Co-IP) assay

BSCs transduced with lentiviruses carrying NS2-flag or mutNS2-flag were lysed in 1X cell lysis buffer (Cat#9803S, Cell Signaling Technology) with protease inhibitor (Cat#11697698001, Sigma-Aldrich) and phosphatase inhibitor cocktails (Cat#524627-1EA, EMD Millipore). After centrifugation to remove debris, the cell lysate was incubated with an anti-FLAG antibody (1:50, Cat#F1804, Sigma-Aldrich) on a rotator overnight at 4°C and then Protein G agarose beads (30 μL, Cat##37478, Cell Signaling Technology) for 3 hours at 4°C. After centrifugation and separation, the supernatant and the pulldown pellet were collected separately. The pulldown pellet was washed with saline five times before Western blot assay. A “conformation Specific” secondary antibody (Cat#5127, Cell signaling technology) was utilized to minimize the detection of unspecific IgG heavy and light chains of the anti-FLAG antibody.

### Quantitative Real Time PCR (qPCR)

Total RNA was extracted from 24-well ALI cultures using the RNeasy Kit (Qiagen, Cat#74106) and then reverse transcribed with SuperScript III Reverse Transcriptase (Thermo Fisher Scientific, Cat#18080-044). Quantitative real-time PCR was performed with SYBR Green Mix (Thermo Fisher Scientific, Cat#4367659) on a CFX96 Real-Time System (Bio-Rad). Relative gene expression was calculated using the ΔΔCt method with GAPDH as the internal control. Primer sequence details are provided and listed in the primer table (Supplementary Data 2).

### Western Blot Assay

ALI protein of 24-well inserts was harvested in 1X cell lysis buffer (Cat#9803S, Cell Signaling Technology) with protease inhibitor (Cat#11697698001, Sigma-Aldrich) and phosphatase inhibitor cocktails (Cat#524627-1EA, EMD Millipore) followed by standard Western blot assays. Primary antibodies include rabbit anti-phosphorylated STAT3 at Tyr705 (1:1000, Cat#9145S, Cell Signaling Technology), rabbit anti-STAT3 (1:1000, Cat#12640S, Cell Signaling Technology), rabbit anti-STAT3 (1:1000, Cat#12640S, Cell Signaling Technology), rabbit anti-H3 (1:1000, Cat#9715, Cell Signaling Technology), rabbit anti-H3K27me3 (1:1000, Cat#9733, Cell Signaling Technology), rabbit anti-H3K27ac (1:1000, Cat#8173, Cell Signaling Technology), rabbit anti-H3K4me3 (1:1000, Cat#9727, Cell Signaling Technology), rabbit anti-RIG-I (1:1000, Cat#3743, Cell Signaling Technology), rabbit anti-Flag (1:1000, Cat#F1804, Sigma-Aldrich), rabbit anti-Caspase-3 (1:1000, Cat#9662, Cell Signaling Technology), rabbit anti-Cleaved Caspase-3 (Asp175) (1:1000, Cat#9661S, Cell Signaling Technology), and mouse anti-β-Actin (1:2000, Cat#A5441, Sigma Aldrich). HRP-conjugated secondary antibodies include goat anti-rabbit (1:3000; Santa Cruz Biotechnology, sc-2004) and goat anti-mouse (1:3000; BD Biosciences, 554002). The antigen-antibody complex was detected by SuperSignal™ West Dura Extended Duration Chemiluminescent Substrate (Cat#34075, Thermo Fisher). Primary antibodies and secondary antibodies were listed in the resource table (Supplementary Data 1)

## QUANTIFICATION AND STATISTICAL ANALYSIS

### Imaging and quantification

Stained slides were imaged using a Nikon Ti inverted fluorescence/confocal microscope or a Nikon DS-Fi2 digital camera microscope. For ALI and lung sections, 4-6 randomly selected non-overlapping 20× images per sample were acquired. Airway epithelial cells were identified by nuclear staining (Hoechst or hematoxylin), and those positive for specific markers were counted and expressed as mean fluorescence intensity (MFI) or a percentage of total epithelial cells. Quantification was performed in a double blinded manner.

### Statistical analysis

Details on statistical analyses, sample sizes, and experimental replicates are provided in the corresponding figure legends, results section, or supplementary section. Statistical comparisons between two groups were performed using unpaired Student’s t-tests. For other comparisons among more than two groups, one-way ANOVA with Dunnett’s multiple-comparison test or two-way ANOVA with Sidak’s multiple-comparison test was applied, as appropriate. All analyses were conducted using GraphPad Prism 8 and p < 0.05 was considered statistically significant.

---

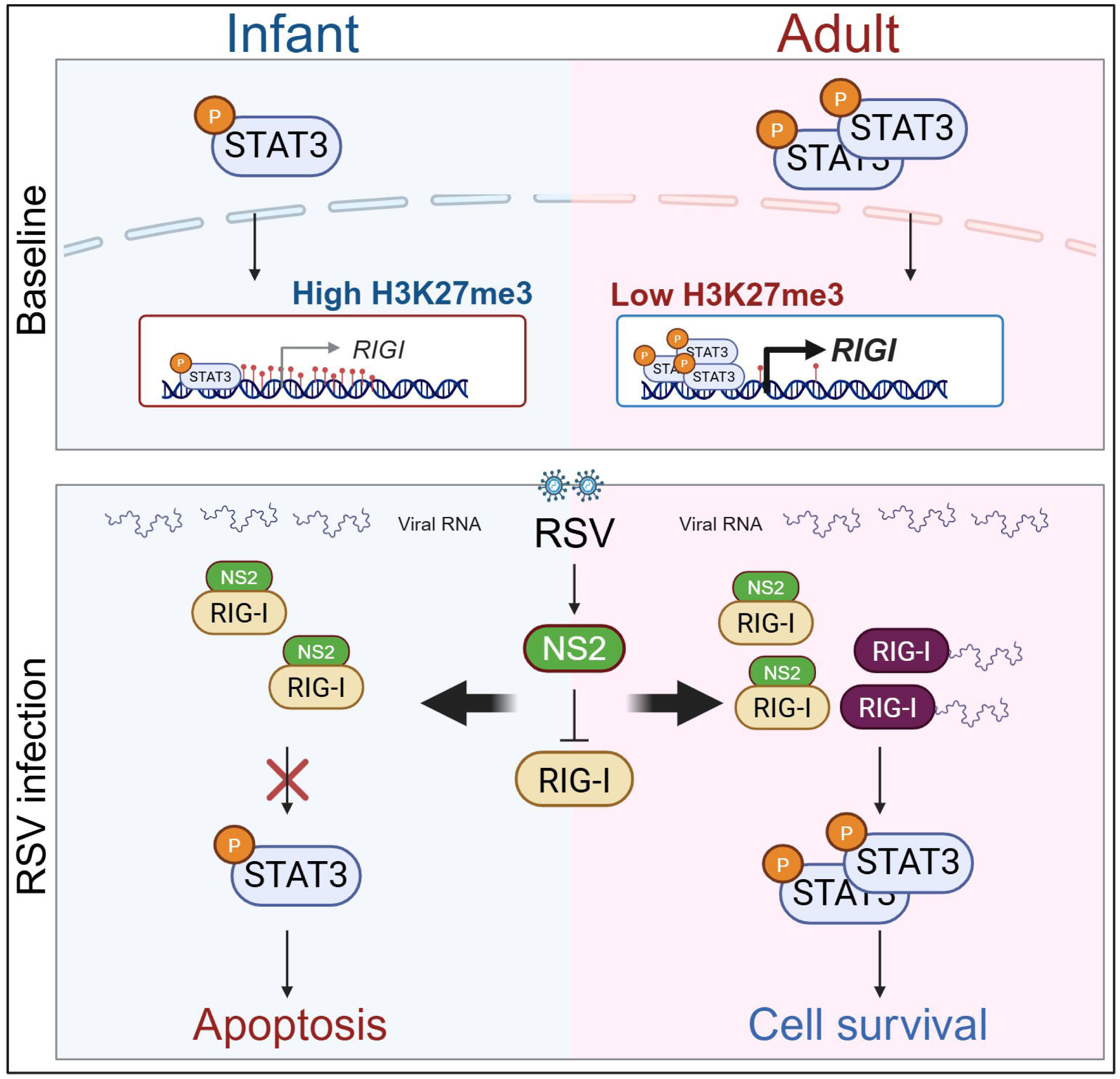

