## Supplementary material for "An immune-poised state of human bronchial epithelial cells mediates RSV resistance in adults": All supplemental files

**SUPPLEMENTARY INFORMATION**

**Supplementary Data 1**: Key resource table.

| REAGENT or RESOURCE | SOURCE | IDENTIFIER |
|---|---|---|
| **Antibodies** | | |
| Mouse anti-RSV Fusion protein | Bio-Rad | Cat# MCA490, RRID: AB_2231368 |
| Mouse anti-p63 [4A4] | Abcam | Cat# ab735, RRID: AB_305870 |
| Rabbit anti-Cleaved Caspase-3 (Asp175) | Cell Signaling Technology | Cat# 9661, RRID: AB_2341188 |
| Rabbit anti-Caspase-3 (Asp175) | Cell Signaling Technology | Cat# 9662, RRID: AB_331439 |
| Rabbit anti-phospho-Stat3 (Tyr705) | Cell Signaling Technology | Cat# 9145, RRID: AB_2491009 |
| Rabbit anti-STAT3 | Cell Signaling Technology | Cat# 12640, RRID: AB_2629499 |
| Mouse anti-FLAG | Sigma-Aldrich | Cat# F1804, RRID: AB_262044 |
| Mouse anti-β-Actin | Sigma-Aldrich | Cat# A5441; RRID: AB_476744 |
| Rabbit anti-Rig-I (D14G6) | Cell Signaling Technology | Cat# 3743, RRID: AB_2269233 |
| Rabbit anti-H3K27me3 | Cell Signaling Technology | Cat# 9733, RRID: AB_2616029 |
| Rabbit anti-H3K27ac | Cell Signaling Technology | Cat# 8173, RRID: AB_10949503 |
| Rabbit anti-H3K4me3 | Cell Signaling Technology | Cat# 9727, RRID: AB_561095 |
| Rabbit anti-H3 | Cell Signaling Technology | Cat# 9715, RRID: AB_331563 |
| Goat anti-rabbit IgG (H+L) AF594 | Thermo Fisher Scientific | Cat# A-11037, RRID: AB_2534095 |
| Goat anti-mouse IgG (H+L) AF488 | Thermo Fisher Scientific | Cat# A-11029, RRID: AB_2534088 |
| Goat polyclonal anti-rabbit, HRP | Santa Cruz Biotechnology | Cat# sc-2004; RRID: AB_631746 |
| Goat polyclonal anti-mouse, HRP | BD Biosciences | Cat# 554002; RRID: AB_395198 |
| **Virus strains and vectors** | | |
| RSV A2 | ATCC | VR-1540 |
| RSV A2-GFP | ViraTree | https://www.viratree.com/ |
| pLJM1-GFP | Addgene | Cat# 19319 |
| pGL3-basic-GFP | Addgene | Cat# 128053 |
| **Cell lines** | | |
| Hep-2 cell line | ATCC | CCL-23 |
| A549 cell line | ATCC | CCL-185 |
| **Chemicals and recombinant proteins** | | |
| Stattic | Tocris | Cat# 2798 |

| | | |
|---|---|---|
| Poly(I:C) (LMW) / LyoVec™ - RIG-I/MDA-5 Agonist | InvivoGen | Cat# tlrl-picwlv |
| RIG012 | MedChem Express | Cat# HY-147124 |
| IL6 (human) Recombinant Protein | Thermo Scientific | Cat# RP8619 |
| Hoechst dye | Thermo Fisher Scientific | Cat# H3570 |
| Y-27632 2HCl | Selleck Chemicals | Cat# S1049 |
| Rapamycin 99% | Fisher Scientific | Cat# R-5000 |
| CHIR 99021 | Tocris Bioscience | Cat# 4423 |
| A83-01 | Tocris Bioscience | Cat# 2939 |
| Doxycycline hyclate, ≥98% (HPLC) | Sigma-Aldrich | Cat# D9891-25G |
| Lipofectamine 3000 Transfection Reagent | Thermo Scientific | Cat# L3000015 |
| Normocin | InvivoGen | Cat# ant-nr-1 |
| Cell Lysis Buffer | Cell Signaling Technology | Cat# 9803S |
| Fluromount-G | Southern Biotech | Cat# 0100-01 |
| Phosphatase inhibitor cocktails | EMD Millipore | Cat# 524627-1EA |
| Protease inhibitor | Sigma-Aldrich | Cat# 11697698001 |
| **Critical Commercial Assays** | | |
| CUT&RUN Assay Kit | Cell Signaling Technology | Cat# 86652 |
| DNA Purification Buffers and Spin Columns (ChIP, CUT&RUN, CUT&Tag) | Cell Signaling Technology | Cat# 14209 |
| DNeasy kit | QIAGEN | Cat# 69504 |
| Standard ABC kit | Vector Laboratories | Cat#PK-6100 |
| Hematoxylin QS | Vector Laboratories | Cat#H-3404 |
| DAB Peroxidase Substrate Kit | Vector Laboratories | Cat# SK-4100 |
| RNeasy kit | QIAGEN | Cat# 74106 |
| Superscript III Reverse Transcriptase Kit | Thermo Fisher | Cat# 18080-044 |
| SYBR Green PCR Master Mix | Thermo Fisher Scientific | Cat# 4367659 |
| SuperSignal West Dura Extended Duration Substrate | Thermo Fisher Scientific | Cat# 34075 |
| Small Airway Epithelial Cell Growth Medium (SAGM) | PromoCell | Cat# C-21070 |
| PneumaCult-ALI Medium | STEMCELL Technologies | Cat# 05001 |
| **Deposited data** | | |

| | | |
|---|---|---|
| Bulk RNA-seq datasets of adult TA BSCs | Bankoti et al. [59] and this paper | GSE212412, GSE242397 |
| Bulk RNA-seq dataset of neonatal TA BSCs | Wagner et al. [22] and this paper | GSE211790, GSE242397 |
| **Software and Algorithms** | | |
| GraphPad Prism 8 | GraphPad Software | https://www.graphpad.com |
| ImageJ 1.49v | ImageJ | https://imagej.nih.gov |
| NIS-Elements (D4.30.01) | NIS-Elements | https://www.microscope.healthcare.nikon.com |
| R (1.30.0) | R Project | https://www.r-project.org |

**Supplementary Data 2.** List of primers and oligos.

| **RT-qPCR primer** | | |
|---|---|---|
| **Gene** | **Forward Primer** | **Reverse Primer** |
| *RSV L2* | GAACTCAGTGTAGGTAGAATGTTTGCA | TTCAGCTATCATTTTCTCTGCCAAT |
| *RSV NS1* | ATGGGCAGCAATTCATTGAG | TGGCATTGTTGTGAAATTGG |
| *RSV NS2* | TTGATGAAAGACAGGCCACA | TGCCAATGCATTCTAAGAACC |
| *CXCL10* | GAAATTATTCCTGCAAGCCAATTT | TCACCCTTCTTTTTCATTGTAGCA |
| *IFNL1* | AATTGGGACCTGAGGCTTCTC | CCAGCGGACTCCTTTTTGG |
| *IFNL3* | TAAGAGGGCCAAAGATGCCTT | CTGGTCCAAGACATCCCCC |
| *ISG15* | CGCAGATCACCCAGAAGATCG | TTCGTCGCATTTGTCCACCA |
| *IFNB1* | ATGACCAACAAGTGTCTCCTCC | GGAATCCAAGCAAGTTGTAGCTC |
| *RIGI* | CTGGACCCTACCTACATCCTG | GGCATCCAAAAAGCCACGG |
| *GAPDH* | GGAGCGAGATCCCTCCAAAAT | GGCTGTTGTCATACTTCTCATGG |
| **shRNA target sequence** | | |
| *RIGI* | GCAAGCCTTCCAGGATTATAT; CCCAATACAAATGGATTATTT | |
| *NS1* | TTGTGTTTGTGCATGTTATTA; GATTCAACAATGACCAATTAT | |
| *NS2* | CAAAGTAGGAAGCACTAAATA; ACAAAGCATACTCCCATAATA | |
| **CUT&RUN primers** | | |
| *RIGI* regulatory primer | ATGACCACCGAGCAGCGA | TGTAGGTAGGGTCCAGGGTC |
| **Oligos for cloning *RIGI* regulatory sequence with STAT3 binding motifs** | | |
| *RIGI*-reg | ACGCAGCCTGCAAGCCTTCCAGGATTATATCCGGAAGACCCTGGACCCTAC | AGCTGTAGGGTCCAGGGTCTTCCGGATATAATCCTGGAAGGCTTGCAGGCTGCGTGTAC |
| mut*RIGI*-reg | ACGCAGCCTGCAAGCCCCTTGAAGCTACGCTTAAGGGACCCTGGACCCTAC | AGCTGTAGGGTCCAGGGTCCCTTAAGCGTAGCTTCAAGGGGCTTGCAGGCTGCGTGTAC |

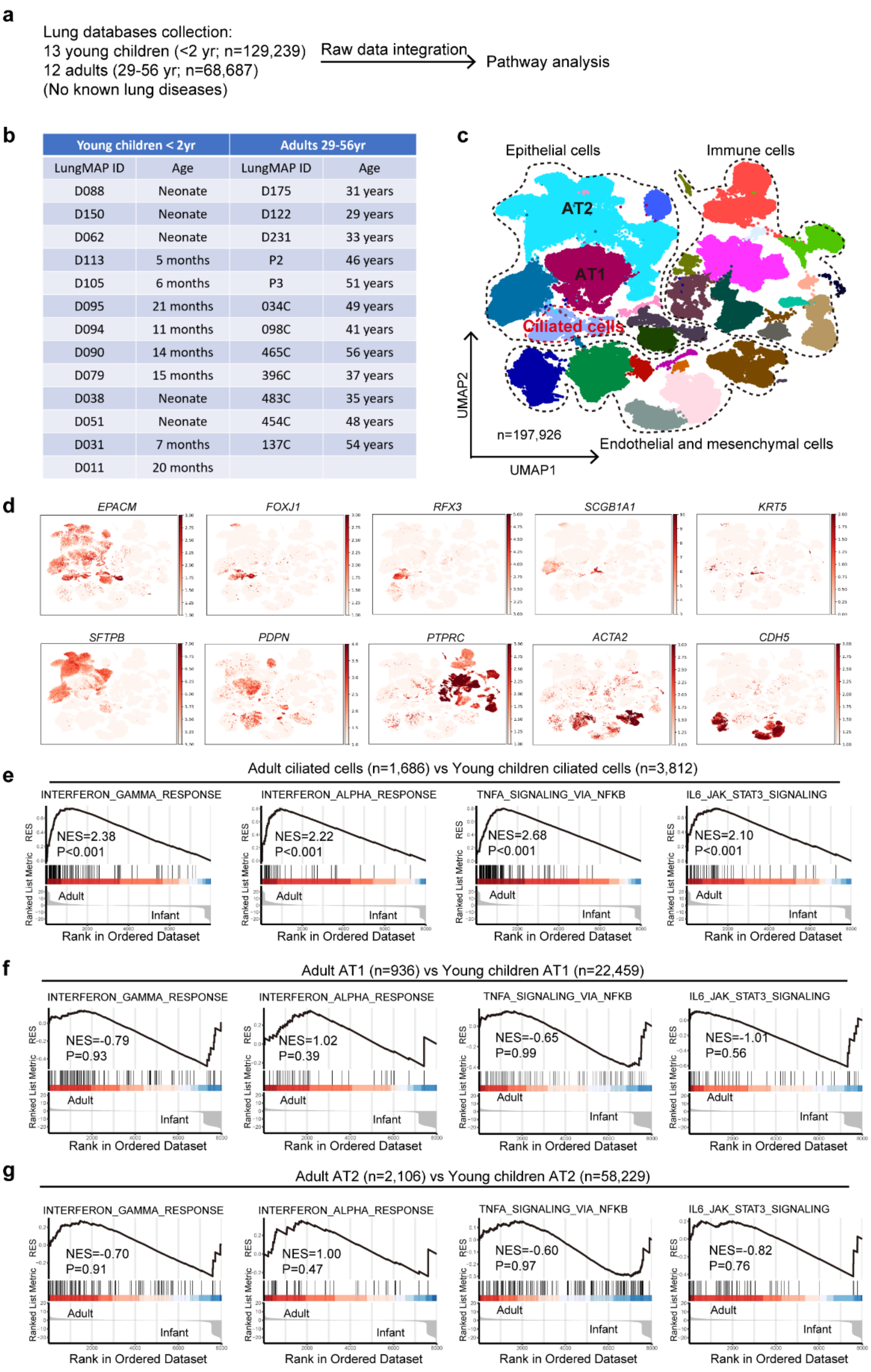


**Extended Data Fig. 1 | Bioinformatic analyses of the scRNA-seq datasets of human lungs available from LungMAP** (**https://www.lungmap.net/**). **a**, Schema of bioinformatic analyses of the scRNA-seq datasets of healthy donor lungs from children younger than 21 months of age (N=13) and adults between 29-56 years of age (N=12). **b**, Donor information. **c**, UMAP plot of scRNA-seq of human donor lungs. **d**, Clusters of different lung cell types with specific cell type markers. **e-g**, Gene set enrichment plots of the IFN, TNFα and IL6 pathways in ciliated cells (**e**), AT1 cells (**f**) and ATII cells (**g**) comparing adults with young children.

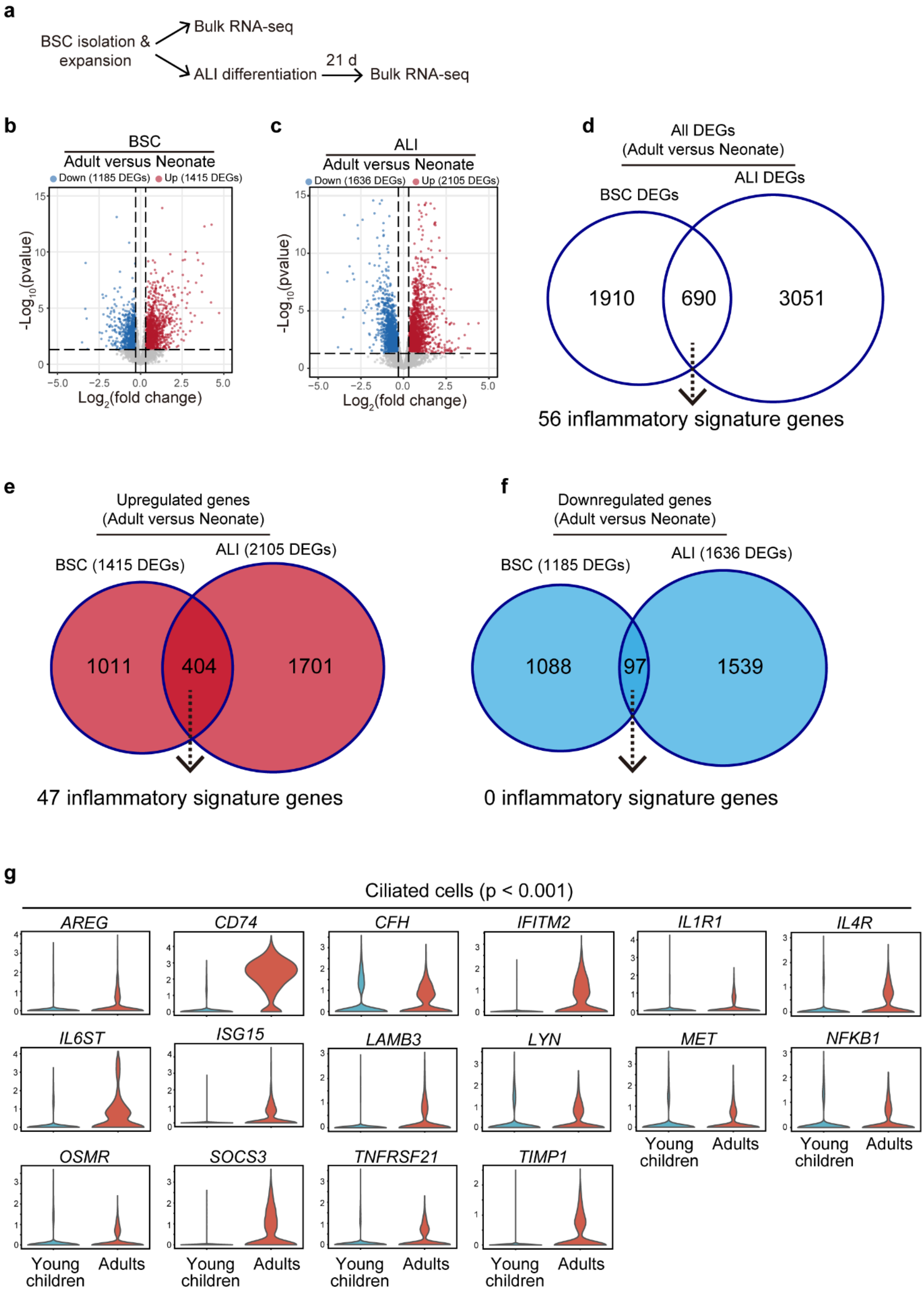


**Extended Data Fig. 2 | Human BSCs and BEpiCs in ALI cultures show age-associated expression of inflammatory response genes.** **a**, Schema of transcriptomic analyses of neonatal and adult BSCs (N=9 per age group) and day 21 ALI cultures (N=3 per age group) by bulk RNA-seq. **b,c**, Volcano plots of differentially expressed genes (DEGs) in BSCs and ALI cultures comparing adults to neonates. The dotted line marks the 30% cutoff. **d**, Venn diagram showing the overlap of age-associated DEGs between BSCs and ALI cultures, including 56 genes within the IFN, TNFα and IL6 response pathways. **e**, Venn diagram showing the overlap of age-upregulated genes between BSCs and ALI cultures, covering 47 genes within the IFN, TNFα and IL6 response pathways. **f**, Ven diagram showing the overlap of age-downregulated genes between BSCs and ALI cultures with 0 genes within the IFN, TNFα and IL6 response pathways. **g**, Violin plots of 16 age-associated genes within the IFN, TNFα and IL6 response pathways that were reliably detected in more than 20% of ciliated cells by scRNA-seq in donor lungs. All 16 genes show elevated levels of expression in adults compared to young children.

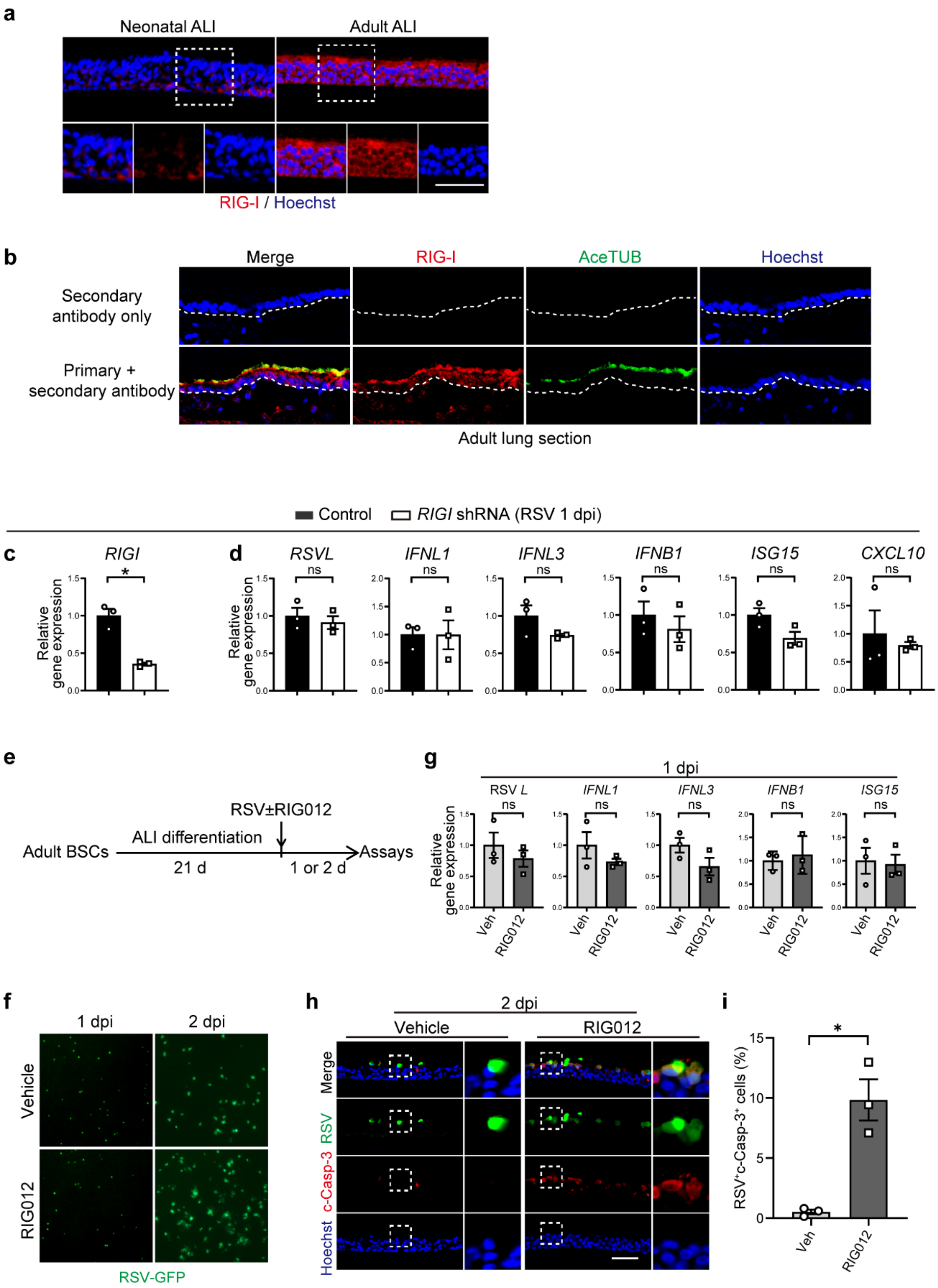


**Extended Data Fig. 3 | Age-associated upregulation of RIG-I in adult BEpiCs is required for RSV resistance**. **a**, Representative images of fluorescence staining for RIG-I in neonatal and adult ALI cultures. The outlined area in the upper panel is enlarged and shown in the bottom panel. RIG-I was detected at a higher level throughout the bronchial epithelium in adult ALI cultures than neonatal ALI cultures. **b**, Representative images of double staining of adult lung sections for AceTUB (a ciliated cell marker) and RIG-I showing co-localization. The dotted line marks the basement membrane of the bronchial epithelium. **c**, Quantification of the knockdown efficiency of the shRNA against *RIGI* in adult BSCs by RT-qPCR. **d**, Relative levels of *RIGI*, RSV *L*, *IFNL1*, *IFNL3, IFNB1*, *ISG15*, and *CXCL10* gene expression in adult BEpiCs in ALI cultures with and without RIG-I knockdown at 1 dpi by RT-qPCR. **e**, Schema of RIG012 (2 μM) treatment of adult BEpiCs in ALI cultures. RIG012 was applied in the bottom chamber at the time of

RSV infection. **f**, Representative images of RSV-GFP-infected adult BEpiCs in ALI cultures with and without RIG012 treatment at 1 dpi and 2 dpi. **g**, Relative levels of RSV *L*, *IFNL1*, *IFNL3*, *IFNB1*, and *ISG15* gene expression with and without RIG012 treatment at 1 dpi by RT-qPCR. **h**, Representative double staining for RSV F and c-Casp-3 in adult BEpiCs using sections of ALI cultures at 2 dpi. **i**, Quantification of the relative abundance of RSV $F^+$c-Casp-$3^+$ cells in infected ALI cultures. Bar graphs show mean ± SEM. *$p<0.05$ and ns, not significant by Student's t-test (two-tailed) in **c**, **d**, **g**, and **i**. Scale bars, 50 μm. Each mark represents one donor. Nuclei were countered stained with Hoechst dye.

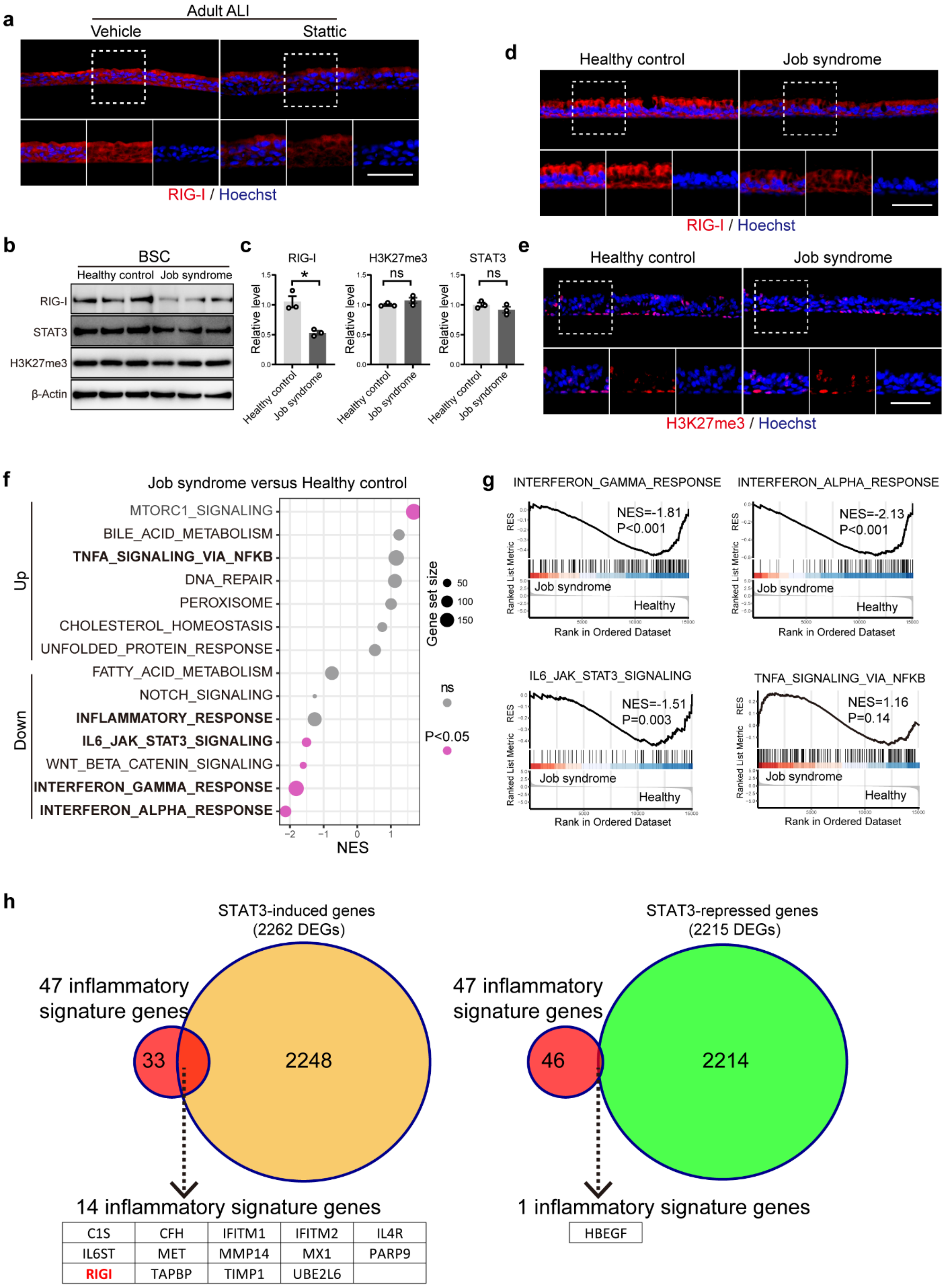


**Extended Data Fig. 4 | RIG-I upregulation in human BEpiCs with age requires active STAT3. a**, Representative images of fluorescence staining for RIG-I in adult BEpiCs in ALI cultures with and without Stattic treatment. The outlined area in the top panel is enlarged and shown in the bottom panels. Stattic (20 μM) was added to the basal medium on day 19 in ALI and reduced the level of RIG-I assayed on day 21. **b**, Western blots of RIG-I, STAT3, and H3K27me3 in adult BSCs from healthy donors (N=3 donors) and a patient with Job syndrome. β-Actin is loading control. **c**, Densitometry of the results in (**b**) normalized to β-Actin. Bar graphs show mean ± SEM. *p<0.05 and ns, not significant by Student's t-test (two-tailed).

**d**, **e**, Representative images of fluorescence staining for RIG-I (**d**) and H3K27me3 (**e**) in day 21 ALI cultures of BSCs of healthy adult donors (N=3 donors) and a Job syndrome patient. The outlined area in the top panel is enlarged and shown in the bottom panels. Job syndrome reduces the level of RIG-I in adult BEpiCs without affecting H3K27me3 in ALI cultures. **f**, Dot plot showing the enrichments of hallmark pathways in Job syndrome BEpiCs in triplicates compared to healthy control BEpiCs in ALI cultures (N=3 donors). **g**, Enrichment plots of the IFN, TNFα and IL6 pathways in the Job syndrome BEpiCs versus healthy control BEpiCs in ALI cultures. **h**, Venn diagrams showing the overlap of the 47 age-upregulated inflammatory response genes with STAT3-regulated genes in BEpiCs in ALI cultures. Among the 15 overlapping genes, 14 genes were induced by STAT3 and one gene was repressed by STAT3. Nuclei in **a**,**d**,**e** were countered stained with Hoechst dye. Scale bars, 50 μm.

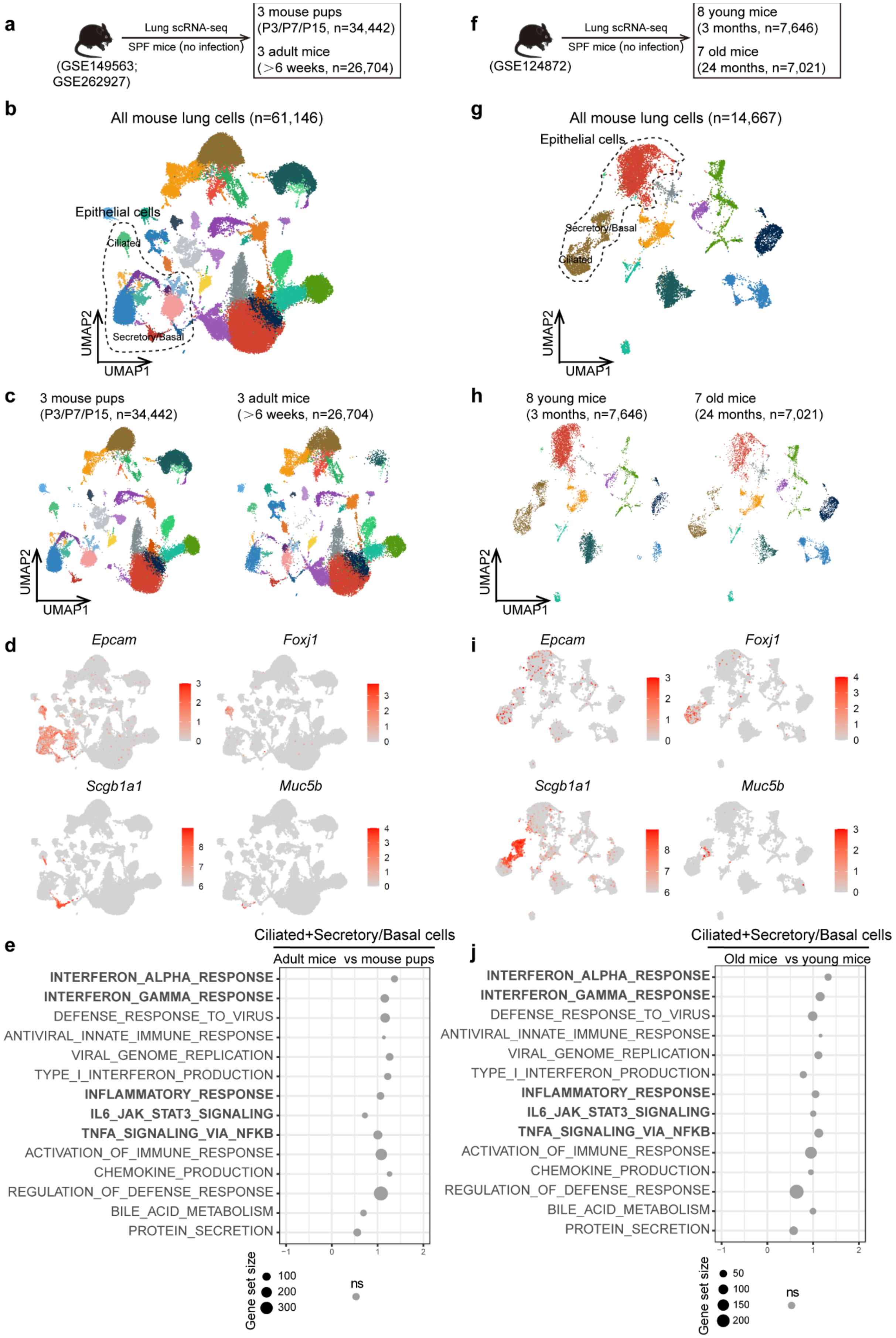


**Extended Data Fig. 5 | Physical age is not sufficient to induce an immune-poised state in adult BEpiCs in SPF mice. a**,**f**, Schema of scRNA-seq of mouse lungs from pups, adults, and aged mice in available datasets (GSE149563, GSE262927, GSE124872). **b**,**g**, Combined UMAP plots showing lung cells of pups and adult mice (N=3 per group) (**b**) and adult and aged mice (**g**). **c**,**h**, UMAP plots showing lung cells in pups and adult mice in (**c**) and adult and aged mice in (**h**). **d**,**i**, Feature plots showing different cell markers in different cell clusters. **e**,**j**, Dot plot showing the enrichments in hallmark pathways in BEpiCs comparing adult mice to mouse pups in (**e**) and adult to aged mice in (**j**).

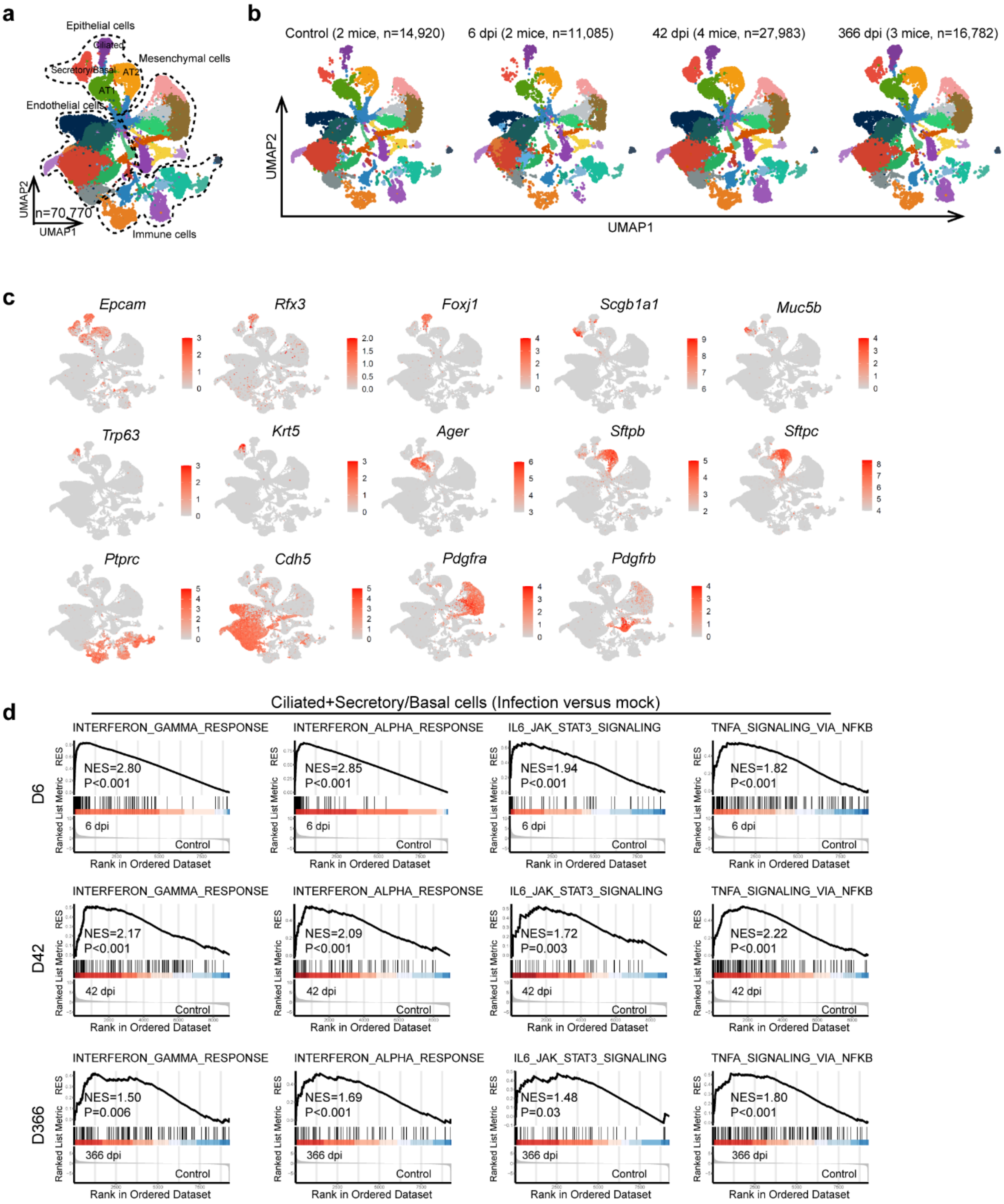


**Extended Data Fig. 6 | Acute respiratory viral infection induces persistent enrichments of inflammatory response pathways in mouse bronchial epithelial cells**. **a**, Combined UMAP plot showing all lung cells (n = 70,770). **b**, UMAP plots showing lung cells of four experimental groups (mock, 6 dpi, 42 dpi, and 366 dpi). **c**, Feature plots showing cell markers in different cell clusters. **d**, Enrichment plots showing the IFN, TNFα and IL6 pathways in mouse bronchial epithelial cells at each time point after infection compared to mock control.

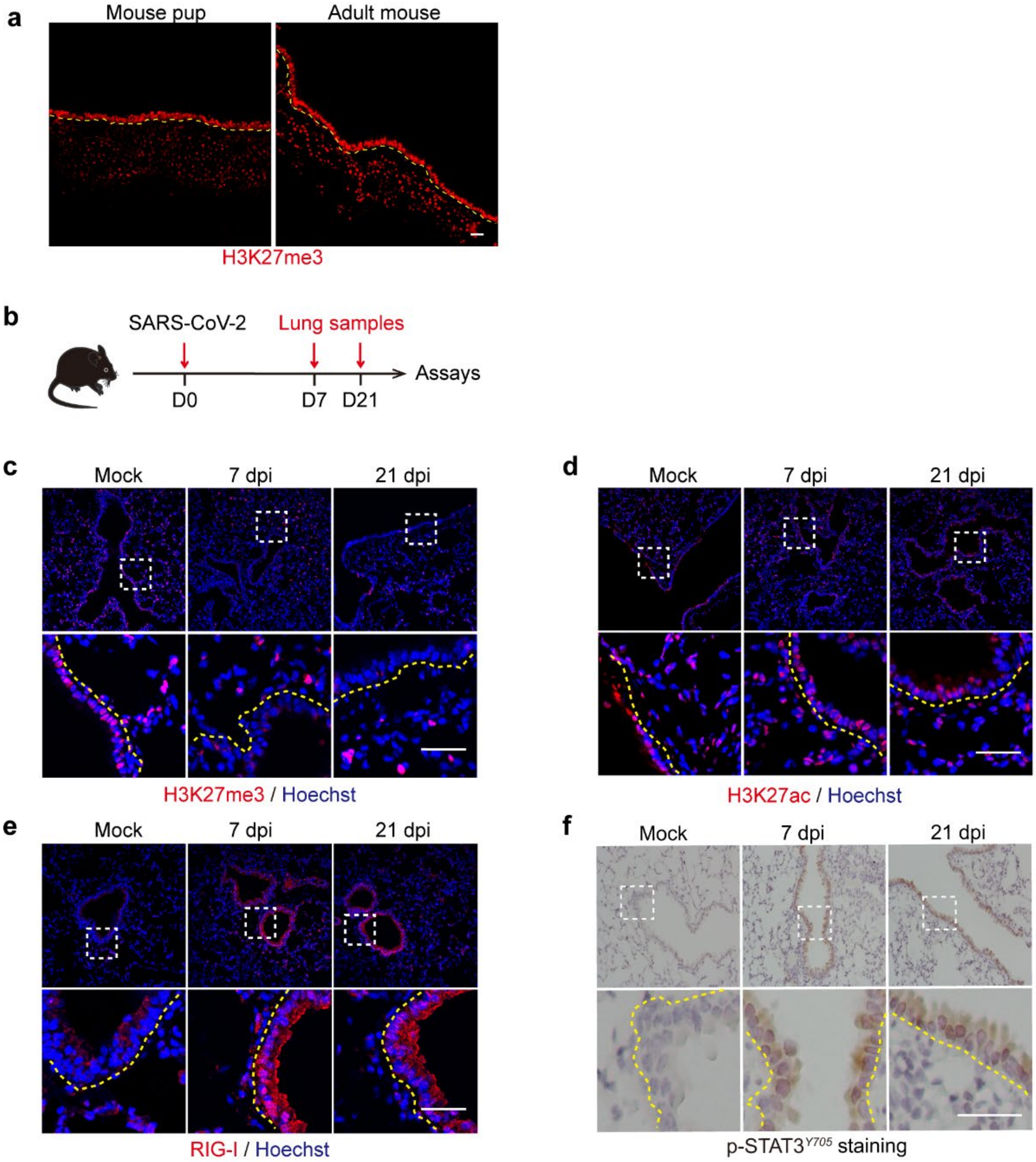


**Extended Data Fig. 7 | Acute SARS-CoV-2 infection induces changes in H3K27me3, RIG-I and p-Stat3$^{Y705}$ in mouse BEpiCs**. **a**, Representative images of fluorescence staining for H3K27me3 in lung sections from mouse pups and adult mice. Comparable and robust levels of H3K27me3 were found in BEpiCs independent of age. **b**, Schema of SARS-CoV-2 infection in adult mice followed by lung harvest and antibody staining. **c-f**, Representative images of antibody staining for H3K27me3 (fluorescence) (**c**), H3K27ac (fluorescence) (**d**), RIG-I (fluorescence) (**e**), and p-STAT3$^{Y705}$ (chromogenic) (**f**) of mouse lung sections. The outlined area in the upper panel was enlarged and shown in the bottom panel. The yellow dotted line marks basement membrane. Cells in the lung mesenchyme showed comparable levels of staining for each marker under the staining and imaging conditions. Nuclei in **c-e** were countered stained with Hoechst dye. Nuclei in **f** were countered stained with hematoxylin. Scale bars, 50 μm.

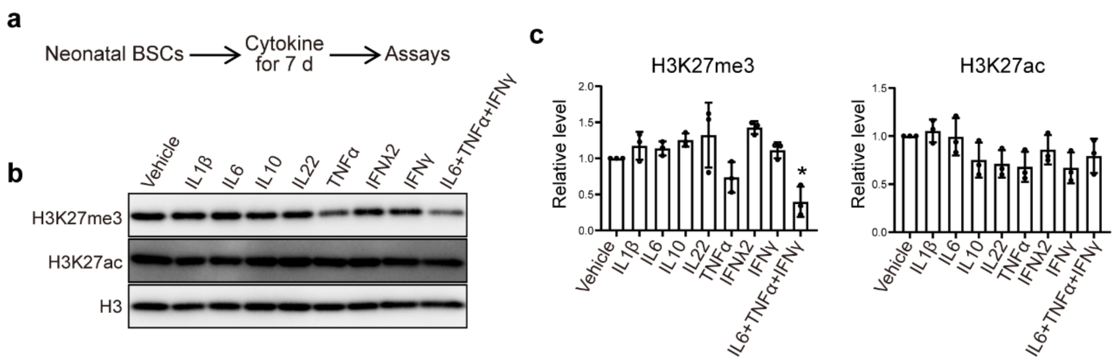


**Extended Data Fig. 8 | Inflammatory cytokine cocktail treatment induces the reduction in H3K27me3 in neonatal BSCs. a**, Schema of neonatal BSCs with cytokine or cytokine cocktail treatment (2 ng/mL for each) for 7 days. **b**, Representative Western blots of H3K27me3 and H3K27ac in neonatal BSCs with and without cytokine treatment. H3 was loading control. **c**, Densitometry of the results normalized to H3. N=3 donors. Bar graphs show mean ± SEM. *$p<0.05$ by one-way ANOVA followed Tukey's multiple comparison test.
